# LENS: Landscape of Effective Neoantigens Software

**DOI:** 10.1101/2022.04.01.486738

**Authors:** Steven P. Vensko, Kelly Olsen, Dante Bortone, Christof C. Smith, Shengjie Chai, Wolfgang Beckabir, Misha Fini, Othmane Jadi, Alex Rubinsteyn, Benjamin G. Vincent

**Affiliations:** Lineberger Comprehensive Cancer Center, University of North Carolina - Chapel Hill, Chapel Hill, NC, USA; Department of Microbiology and Immunology, University of North Carolina - Chapel Hill, Chapel Hill, NC, USA; Department of Medicine, Brigham and Women’s Hospital, Boston, MA, USA; Uber Technologies, Inc., San Francisco, CA, USA; Department of Genetics, University of North Carolina - Chapel Hill, Chapel Hill, NC, USA; Curriculum in Bioinformatics and Computational Biology, University of North Carolina - Chapel Hill, Chapel Hill, NC, USA; Computational Medicine Program, University of North Carolina - Chapel Hill, Chapel Hill, NC, USA; Department of Medicine, Division of Hematology, University of North Carolina - Chapel Hill, Chapel Hill, NC, USA

## Abstract

**Motivation:** Elimination of cancer cells by T cells is a critical mechanism of anti-tumor immunity and cancer immunotherapy response. T cells recognize cancer cells by engagement of T cell receptors with peptide epitopes presented by major histocompatibility complex (MHC) molecules on the cancer cell surface. Peptide epitopes can be derived from antigen proteins coded for by multiple genomic sources. Bioinformatics tools used to identify tumor-specific epitopes via analysis of DNA and RNA sequencing data have largely focused on epitopes derived from somatic variants, though a smaller number have evaluated potential antigens from other genomic sources.

**Results:** We report here an open-source workflow utilizing the Nextflow DSL2 workflow manager, Landscape of Effective Neoantigen Software (LENS), which predicts tumor-specific and tumor-associated antigens from single nucleotide variants, insertions and deletions, fusion events, splice variants, cancer testis antigens, overexpressed self-antigens, viruses, and endogenous retroviruses. The primary advantage of LENS is that it expands the breadth of genomic sources of discoverable tumor antigens using genomics data. Other advantages include modularity, extensibility, ease of use, and harmonization of relative expression level and immunogenicity prediction across multiple genomic sources. We present an analysis of 115 acute myeloid leukemia (AML) samples to demonstrate the utility of LENS. We expect LENS will be a valuable platform and resource for T cell epitope discovery bioinformatics, especially in cancers with few somatic variants where tumor-specific epitopes from alternative genomic sources are an elevated priority.

**Availability:** More information about LENS, including workflow documentation and instructions, can be found at https://gitlab.com/landscape-of-effective-neoantigens-software

**Supplementary information:** Supplementary data are available at *Bioinformatics* online.

## 1 Introduction

Tumor-specific and tumor-associated antigens are of great interest for understanding cancer immunobiology and developing personalized immunotherapy approaches including vaccination or adoptive T cell therapy. Unfortunately, predicting which tumor antigens are immunogenic *in vivo* is challenging (Wells *et al*., 2020; Smith *et al*., 2019c). These predictions are complicated by several factors influencing the appropriateness of candidate tumor antigens. Specifically, only a subset of tumor-specific or tumor-associated variants or sequences within a patient will be transcribed, translated, and processed by the proteosome. Only a subset of peptides generated through protein degradation will be presented on the cell surface by MHC and only a further subset of those will result in T-cell recognition, activation and cytotoxicity. Despite recent advances in understanding peptide attributes associated with *in vivo* immunogenicity, other factors including RNA editing, post-translational modifications, and peptide splicing may influence the effectiveness of computational predictions (Liepe *et al*., 2018; Zhang *et al*., 2018; Rolfs *et al*., 2018). Improved cancer antigen prediction using genomics data could empower more detailed studies of anti-tumor T cell responses as well as better personalized combination immunotherapy strategies (Hu *et al*., 2021).

Multiple workflows exist to predict tumor antigens from high throughput sequencing data. These include OpenVax, pVACTools, and nextNEOpi among others (Kodysh and Rubinsteyn, 2020; Hundal *et al*., 2020; Rieder *et al*., 2022). These tools allow users to study associations between tumor antigen burden and immunotherapy response, evaluate tumor-antigen specific T cell responses, and support clinical trials of personalized immunotherapy targeting predicted tumor antigens (Litchfield *et al*., 2021; Kodysh and Rubinsteyn, 2020; Ott *et al*., 2017; Sahin *et al*., 2017; Lowery *et al*., 2022; Caushi *et al*., 2021; Hu *et al*., 2021). Here we present an extensible, modular, and open-source workflow, Landscape of Neoantigen Software (LENS), coupled with a Nextflow-based analysis platform, Reproducible Analyses Framework and Tools (RAFT, see Section 2.1), which addresses shortcomings of current neoantigen workflows, expands the repertoire of predicted tumor antigens, and serves as a springboard towards community-driven advances in neoantigen prediction.

## 2 Landscape of Effective Neoantigens Software

Landscape of Effective Neoantigens Software (LENS) is a modular, extensible workflow which predicts tumor antigens from an array of sources including somatic single nucleotide variants (SNVs), conservative, disruptive, and frameshift insertions and deletions (InDels), splice variants, in-frame and frameshift fusion events, viruses, endogenous retroviruses, and cancer testis antigens/self-antigens (Figure 1). LENS includes phasing and germline variant information in epitope identification, and it harmonizes variant RNA expression across genomic sources to provide a more usable relative expression ranking each peptide epitope. More information about LENS, including workflow documentation and instructions for running LENS can be found at https://gitlab.com/landscape-of-effective-neoantigens-software.

**Fig. 1:**
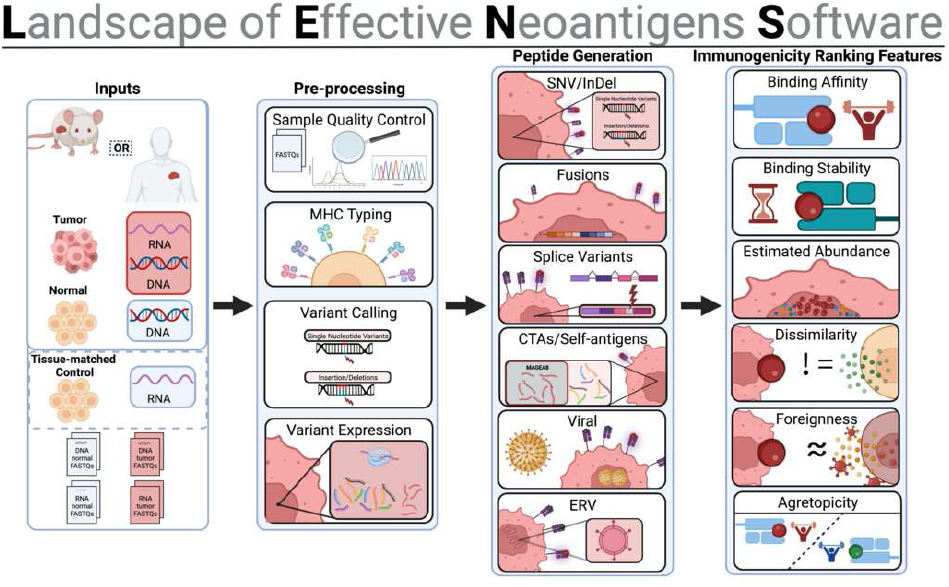
Landscape of Effective Neoantigens Software (LENS): A modular workflow for predicting Class I neoepitopes from SNVs, InDels, fusion events, splice variants, CTAs, self-antigens, viruses, and ERVs.

Here we provide an overview of the LENS workflow, describe its advantages over currently available workflows, describe its technical aspects, and discuss the results of running LENS on 115 Acute Myeloid Leukemia patients.

### 2.1 Implementation

The framework running LENS is a Nextflow wrapper called RAFT (Reproducible Analyses Framework and Tools) which enables our goals of modularity, extensibility, and improved support for collaboration (Di Tommaso *et al*., 2017). RAFT supplements the Nextflow DSL2 with a collection of tool-level modules, workflow modules, and automation to ease workflow creation and running (e.g. module dependency resolution, parameter centralization, etc.). A full description of RAFT’s capabilities can be found at RAFT’s GitHub: https://gitlab.com/landscape-of-effective-neoantigens-software/raft.

### 2.2 Flexibility

The modular design of LENS allows high flexibility in its execution. This presents itself in many forms including user-definable references files (FASTAs, GTF/GFFs, BED files, etc.) and parameters on either global or workflow-specific scopes. As a result, users maintain fine-grain control over workflow and tool-level behaviors. This allows manipulation of workflows to account for variability in sample type (e.g. fresh frozen vs. formalin fixed) or changing filtering criteria. Perhaps most importantly, the modular nature of LENS allows users to introduce novel tools or replace current tools with a better suited one. There may be other scenarios where swapping tools may be desirable or necessary as there are a variety of variant callers, fusion callers, and copy number inference tools available with their individual benefits and disadvantages (Haas *et al*., 2019). Replacing the LENS default tools with these alternatives or defining new ensemble approaches may produce richer and more meaningful results. Additionally, users may want to change tools to be within compliance of any license requirements encompassing their usage.

### 2.3 Extensibility

Properly leveraging novel technologies, protocols, and public datasets is crucial for progress towards meaningful and impactful insights in tumor antigen discovery. In light of this, we designed LENS such that it can be augmented through creation of new modules and workflows. LENS reports can further be supplemented by including published pMHC-specific empirical immunogenicity data as more become available. Inclusion of single-cell sequencing modules would help disentangle the genetic and transcriptional heterogeneity within a patient’s tumor that cannot be observed in bulk sequencing data. Long read sequencing allows for detection of large structural variants and provides improved resolutions of haplotypes near tumor peptide-generating sequences of interest. Other technologies, such as ribosome profiling (Ribo-seq), will not only improve current tumor antigen predictions by confirming translation reading frames, but can open new tumor antigen sources such as “genomic dark matter” arising from non-canonical reading frames (Delgado *et al*., 2014). We consider the version of LENS presented here to be a snapshot of the continuously improving workflow that will aid the immuno-oncology community towards improved therapeutics.

### 2.4 Comparison to Other Workflows

Neoantigen workflows have become increasingly popular as the value of personalized immunotherapy has been realized. There are currently dozens of neoantigen workflows publicly available including pVACtools, the OpenVax workflow, nextNEOpi, and MuPeXi (Hundal *et al*., 2020; Kodysh and Rubinsteyn, 2020; Rieder *et al*., 2022; Bjerregaard *et al*., 2017). A comprehensive comparison among all available workflows is available in the literature (Rieder *et al*., 2022). LENS improves upon previous offerings by providing an end-to-end solution utilizing containerized tools and modular workflows, expanding the types of tumor antigens predicted (to include splice variants, cancer-testis antigens/self-antigens, oncogenic viruses, and endogenous retroviruses), providing a harmonized tumor antigen abundance quantifier, and allowing customization of the workflow. A feature-level comparison against competing workflows that have been used in clinical trials can be found in Supplemental Table 1. A high-level comparative summary between LENS and twenty-six other workflows can be found in Supplemental Table 4.

### 2.5 Workflow Overview

The LENS workflow orchestrates over two dozen separate tools to generate tumor antigen predictions. LENS currently supports tumor antigen detection from the following tumor antigen sources: SNVs, InDels, fusion events, splice variants, viruses, endogenous retroviruses, cancer testis antigens, and self-antigens. LENS will be expanded to include support for single-cell data, long read data, external reference data, and additional bioinformatics tools. A brief summary on peptide generation approaches for each workflow can be found in Supplemental Table 4.

### 2.6 Data Pre-Processing, Read Alignment, and Transcript Quantification

LENS processes and aligns input FASTQS against a reference genome with an annotation file. Alignment sanitization is performed to ensure high quality inputs for downstream tumor antigen source-specific workflows. Transcripts defined within the GTF annotation file are then quantified. The pre-processing workflow results in numerous intermediate files required for each tumor antigen workflow and is visualized in Supplemental Figure 1.

### 2.7 Single Nucleotide Variants and Insertions/Deletions

Somatic single nucleotide variants (SNVs) and insertion/deletion variants (InDels) may result in targetable tumor-specific peptides. These peptides may be a direct result of a somatic variant or may be a downstream coding consequence of the variant. They should not exist in non-tumor tissues, so T cells that target them could escape negative selection in the thymus, making them attractive targets for antigen-specific immunotherapy (Smith *et al*., 2019a). Vaccines targeting SNV-derived tumor-specific antigens have been used in clinical trials, with strategies including peptide vaccines (Lilleby *et al*., 2017; Obara *et al*., 2017), dendritic cell vaccines (Filley and Dey, 2017; Carreno *et al*., 2015), DNA vaccines (Li *et al*., 2021), and RNA vaccines (Sahin *et al*., 2017).

The LENS SNV and InDel workflows filter somatic variants using a consensus approach. By default, somatic variants are called using three variant callers (Mutect2, Strelka2, and ABRA2), and variants within the intersection of the appropriate callers (MuTect2 and Strelka2 for SNVs; all three callers for InDels) are considered (McKenna *et al*., 2010; Kim *et al*., 2018; Mose *et al*., 2019). Users may also use a union of variants as well as other variant callers such as VarScan2 (Lang *et al*., 2022). Requiring consistent variant detection among callers culls potential false positive variants leaving a set of high confidence calls. The intersected VCF is filtered for coding variants (missense SNVs, conservative and disruptive in-frame InDels, and frameshift InDels) for further processing. Variants are filtered by two criteria at different points in the workflow: 1) their relative transcription abundance must exceed an abundance percentile (e.g. 75%) and 2) sequence capable of coding variant-harboring peptides must be detectable in the patient’s tumor RNA sequencing data. Expression filtering creates a stable set of SNVs/InDels that are likely transcribed and translated and may be presented by the MHC on the tumor cell surface. Accurately generating tumor peptides from the patient’s somatic variants requires phasing candidate somatic variants with neighboring germline variants (Supplemental Figure 2). LENS calls germline variants with DeepVariant and performs phasing using the patient’s DNA- and RNA sequencing data through WhatsHap (Martin *et al*., 2016). Phased heterozygous germline variants within the same haplotype block as a candidate somatic variant are incorporated into the resulting peptide where appropriate. This step ensures the predicted peptides more accurately reflect those potentially presented by the patient’s tumor.

The report generated by the SNV workflow includes several metrics relevant to peptide prioritization (described in Section 2.14), the gene and transcript harboring the expressed variant, the transcript’s relative abundance (in transcripts per million), as well as the number of RNA sequencing reads from the peptide’s genomic origin containing a sequence which translates to the peptide (Jurtz *et al*., 2017; Rasmussen *et al*., 2016; Richman *et al*., 2019). The last metric will be discussed in more detail later, but serves as an estimated proxy for peptide abundance harmonized across tumor antigen workflows. Furthermore, LENS estimates variant cancer cell fraction with PyClone-VI by using copy number alteration (CNA) data from Sequenza along with MuTect2 allele frequencies (Gillis and Roth, 2020; Favero *et al*., 2015). Understanding the resulting distribution of tumor sub-populations and estimating the clonality of variants will allow for improved prioritization of predicted neoantigens. The SNV/InDel workflow is visualized in Supplemental Figure 3.

Many SNV and in-frame InDel neoantigens have shown limited utility for inducing a strong immune response (Wells *et al*., 2020). SNV/InDel neoantigen therapies have focused on tumor types known to have high tumor mutational burden such as melanoma. These therapeutic approaches may not be appropriate for patients with lower tumor mutational burden (TMB) tumors, such as acute myeloid leukemia (AML). Furthermore, SNVs and InDel neoantigens tend to be private to individuals which limits the possibility of “off the shelf” neoantigen vaccines. These considerations suggest other tumor antigen sources beyond somatic variants are worthy of further consideration.

### 2.8 Splice Variants

Aberrant splicing events create tumor-specific transcripts through intron retention, exon skipping, or altered splice targeting (Supplemental Figure 4). These transcripts may be translated and processed into targetable tumor-specific peptides. Ninety-four percent of human genes have intronic regions (Zhang *et al*., 2021) and most undergo alternative splicing to generate transcriptomic and proteomic diversity in order to fulfill a wide variety of protein functions (Frankiw *et al*., 2019). Alternative splicing is influenced by RNA structure, chromatin structure, and transcription rate as well as splice site generating or disrupting cis-acting mutations and trans-acting mutations that alter splicing factors to cause aberrant splice variants in other genes (Smith *et al*., 2019a).

LENS utilizes NeoSplice, a k-mer searching and splice graph traversal algorithm, to detect tumor-specific splice variants and neoantigens derived from them (Chai *et al*., 2022). NeoSplice considers patient-specific splice variants derived from their tumor RNA sample relative to splice variants from a tissue-matched normal RNA sample. Differences in k-mer distributions between the tumor and normal splice variants allow for high confidence detection of tumor-specific splice variants. Peptide abundance is estimated by counting the number of reads containing the complete peptide’s coding sequence mapping to the splice variant genomic origin. The splice variant workflow is visualized in Supplemental Figure 5.

### 2.9 Fusion Events

Scenarios in which a translocation, deletion, or inversion causes two previously genomically distant loci to become neighboring sequences or “fuse” within the genome are also of interest as sources of tumor antigens. This combination of two naturally occurring intragenic coding sequences can create novel peptides that may serve as immunogenic epitopes. This effect may be amplified when fusion events result in translational frameshifts that are as potentially immunogenic as splice variants (Wang *et al*., 2021), theoretically moreso than SNV-derived neoantigens (Yang *et al*., 2019).

LENS utilizes STARFusion for detection of fusion events using recommended default parameters (Haas *et al*., 2017). Similar to the SNV and InDel workflows, homozygous germline variants are incorporated into the fused coding sequence prior to translation using phased germline VCFs. Proteins derived from in-frame fusion events are truncated upstream to a user-specified length and downstream of the fusion junction and processed through a suite of tools to characterize potential pMHCs (see Section 2.14). Frameshift-derived proteins are truncated upstream of the fusion junction and include all downstream sequence until the first stop codon. Peptides are filtered and quantified by checking for their coding sequence within the RNA reads that either map across a junction point or are mapped to both sides of the junction point. The fusion workflow is visualized in Supplemental Figure 6.

### 2.10 Tumor-specific Viruses

Some viruses such as human papillomavirus (HPV), Epstein-Barr virus (EBV), human T cell leukemia/lymphoma virus type 1 (HTLV1), and hepatitis C virus (HCV) are associated with development of cancers in the tissues they infect earning them the label of oncogenic viruses (Zhao *et al*., 2021; Perz *et al*., 2006; Mesri *et al*., 2010). Viruses can drive oncogenesis through disruption of host cell growth and survival, induction of DNA damage response causing host genome instability, by causing chronic inflammation and tissue damage, or by causing immune dysregulation creating a more permissive immune environment for tumorigenesis (Tashiro and Brenner, 2017). Antigens derived from the virus are regarded as tumor-associated antigens, yet are immunologically more foreign compared to self-derived peptides, making them potential immunotherapeutic targets.

LENS detects viral-derived tumor antigens with VirDetect, a previously developed workflow designed to detect viral contamination within RNA sequencing data (Selitsky *et al*., 2020). Specifically, RNA reads are mapped against the reference genome and any reads that cannot be mapped to the reference are diverted to the VirDetect workflow. VirDetect aligns these unmapped reads to coding sequences of over 1,900 vertebrate viruses. Homozygous germline variants detected within the RNA sequencing data through BCFtools are incorporated into the viral sequence prior to translation. Translated viral peptide sequences are considered as potential tumor antigens for downstream processing. Peptides are quantified by counting occurrences of peptide-associated coding sequences within the reads that map to expressed viral coding sequences. The viral workflow is visualized in Supplemental Figure 7.

### 2.11 Endogenous Retroviral Elements

Endogenous retroviruses (ERVs) are, unlike typical viruses, integral to the human genome and make up roughly 8% of the genome (Grandi and Tramontano, 2018). Some retroviral elements have retained intact open reading frames that may be transcribed, translated, processed, and presented by MHCs on the cell surface under abnormal transcriptional regulation within a tumor. Their lack of central tolerance, elevated abundance, and ability to be expressed under chaotic tumor conditions make endogenous retroviruses intriguing potential sources of tumor antigens. Unsurprisingly, ERVs have been a focal point of tumor antigen research due to evidence suggesting CD8+ T cell recognition of ERV-derived peptides results in an immunogenic response (Smith *et al*., 2019b; Bonaventura *et al*., 2022).

LENS detects ERV-derived candidate peptides through use of the gEVE (genome-based Endogenous Viral Element) database which includes computationally predicted retroviral element open reading frames (ORFs) from which antigens may be generated (Nakagawa and Takahashi, 2016). Expressed ORFs have homozygous germline variants integrated into their sequences prior to translation. We assume some ERVs have natural low levels of expression in some normal tissues, so differential expression filtering is used to narrow the considered pool (Supplemental Figure 8). Specifically, tissue-matched normal control samples are processed through an ERV quantification workflow and the resulting raw counts are normalized to patient-specific ERV counts through EdgeR and ERV ORFs which exceed a fold-change threshold (*log*(*CPM*_*tumor*_ + 1) − *log*(*CPM*_*normal*_ + 1) *>* 1, see Supplemental Figure 12) are considered for downstream processing (Robinson *et al*., 2010). ERV peptides are quantified by counting peptide-coding sequences among reads that map to expressed ORFs. The ERV workflow is visualized in Supplemental Figure 9.

### 2.12 Cancer Testis Antigens and Self-Antigens

Tumor self-antigens are an additional potential target for immunotherapy. These antigens are non-mutated self antigens that may be overexpressed within a tumor. These antigens are traditionally viewed as suboptimal therapeutic targets for two reasons: (1) T cells recognizing self antigens would be expected to be deleted in the thymus and (2) autoimmunity is difficult to avoid when targeting antigens expressed in both normal and tumor tissues. However, work has been done to identify methods of self antigen targeting while minimizing autoimmunity. The pool of self antigens to select from for targeting is also large, as any given tumor will present many self antigens, making them potential vaccine targets (Cheever *et al*., 2009; Bright *et al*., 2014). Genes normally expressed in the immune privileged tissues, such as testis tissue, may be transcriptionally active in tumors. Antigens derived from these transcripts are commonly referred to as cancer testis antigens (CTAs) and are of particular interest due to their potential immunogenicity. These antigens are derived from normal genes that are usually expressed either during early development or in adulthood only within immune privileged tissues, but may become overexpressed within tumors such as melanoma, breast cancer, or bladder cancer. This abnormal expression makes them promising targets due to their lack of expression in normal tissues undergoing immune surveillance (Mitchell *et al*., 2021).

LENS accepts a user-provided list of CTA and self-antigen gene identifiers, but defaults to a set of cancer testis genes from CTDatabase (Almeida *et al*., 2009). This list includes genes that are germline-biased and highly expressed in cancers, but have not necessarily been shown to initiate an immune response when targeted. The CTA and self-antigen transcripts are first filtered for transcription abundance exceeding a user-specified percentile threshold. Transcripts passing the filter then have homozygous germline variants incorporated and the resulting predicted peptides are run through a suite of tools to characterize candidate pMHCs. pMHCs with high binding affinity and high pMHC stability are quantified by counting occurrences of peptide-coding sequence observed within reads mapping to the peptide’s genomic origin. The CTA/Self-antigen workflow is visualized in Supplemental Figure 10.

### 2.13 Tumor Antigen Peptide Characterization

Peptides are generated by antigen source-specific workflows and then combined into a single set. These peptides are combined with the patient’s HLA alleles to characterize aspects of the pMHC. Calculated peptide features include estimates of binding affinity, binding stability, proteosomal processing score, presentation score, dissimilarity, foreignness, and agretopicity. Supported tools are currently NetMHCpan (binding affinity), NetMHCstabpan (binding stability), MHCFlurry (binding affinity, antigen processing, and presentation), DeepHLAPan (binding affinity and immunogenicity), and antigen.garnish (dissimilarity and foreignness).

### 2.14 Harmonization of Peptide Abundance Estimates

The tumor antigen workflows within LENS have a variety of quantification metrics generated by the individual workflows. These metrics include Transcripts per Million (TPM) for transcript quantification in the SNV and InDel workflows, Fusion Fragments per Million (FFPM) for fusion quantification in the fusion workflow, minimum observed expression for the splice variant quantification in the splice workflow, and read count for viral expression in the viral workflow. Each metric described may be broadly relevant to tumor antigen quantification, but most have at least two issues: 1) they conflate transcript abundance of the peptide-generating coding sequence with the transcript abundance of its wildtype counterpart and 2) they do not allow meaningful comparisons of peptide abundance among workflows. As an example of the first issue, consider TPM, a metric commonly used to represent relative transcript abundance. TPMs have been used as a filtering criterion for SNV and InDel neoantigens, but they represent the abundance of the transcript (rather than just the peptide-generating sub-sequence) and may include alleles that do not code for the peptide of interest. To address this, we developed a novel quantification strategy utilizing the observed occurrences of the nucleotide sequence responsible for coding the peptide from the peptide’s genomic origin from the patient’s tumor RNA sequencing data (Supplemental Section 4.1). This metric resolves both of the limitations of currently used metrics and allows for prioritization by abundance across multiple tumor antigen types.

### 2.15 Visualization

Neoeptitope confirmation and prioritization are key to selecting potentially therapeutically useful targets for vaccine development. To address this need, we developed LENS Viz, a Shiny web app available at https://lens.shinyapps.io/lensviz (or run locally using code at https://gitlab.com/landscape-of-effective-neoantigens-software/lensviz) (Supplemental Figure 11). LENS Viz allows interactive exploration of the data. This results in an improved understanding of binding affinity, binding stability, and RNA read support on a per-peptide and per-antigen source basis and how modifying thresholds affect the set of potential tumor antigens of interest. It also includes an interactive IGV widget to allow users to view read-based evidence of tumor antigen sources in the tumor samples (and the lack of evidence in the normal sample).

### 2.16 Murine Support

Research using murine models of cancer to better understand anti-tumor immune mechanisms has been crucial to improving human health through immuno-oncology. To this end, LENS was designed to support usage of mouse samples and references to predict potential pMHCs for common research strains. Instructions for running LENS on mouse samples can be found at https://gitlab.com/landscape-of-effective-neoantigens-software/nextflow/modules/tools/lens/-/wikis/Running-LENS. We have tested LENS on the BBN963 strain, a mouse model of bladder cancer, and discovered potential therapeutic targets that are being followed up with experimental testing.

## 3 Results

### 3.1 Applying LENS to TCGA-LAML

The design principles of LENS allow it to avoid constraints around specific tumor types or tumor antigen sources. We demonstrate LENS by processing data from several patients of the widely available TCGA-LAML (Acute Myeloid Leukemia) dataset. Acute myeloid leukemia is suitable for demonstrating LENS due to its relatively low tumor mutational burden (TMB) compared to other tumor types. This low TMB presents a situation where non-SNV- and InDel-derived neoantigens are crucial. The TCGA-LAML dataset also contains sufficient patient-level data including the three sample types (normal exome sequencing, abnormal (tumor) exome sequencing, abnormal (tumor) RNA sequencing) for 115 patients. Here, we discuss the results for each tumor antigen workflow currently available from LENS. This is not intended to be an exhaustive analysis of LENS outputs, but rather to provide an example of its utility in tumor antigen prediction using genomics data.

### 3.2 General Observations among Tumor Antigen Sources

TCGA-LAML patients show a range of predicted tumor antigen counts from 47 to 1,789 (median: 366) (Supplemental Figure 13). Predicted antigen counts among patients vary by genomic source. Specifically, CTA/Self-antigen and ERV peptides made up over 37,000 total predicted peptides among patients (37,349/41,395 (90.2%)) while the remaining sources (SNVs, InDels, fusion events, splice variants, and viruses) consisted of 4,046 peptides (9.8%). This disparity is likely explained by two factors: 1) ERVs are highly abundant within the genome and 2) the entirety of each CTA/Self-antigen’s coding sequence and ERV’s ORF sequence is used for peptide generation. This increase of “raw material” compared to point neoantigen targets, like SNVs and InDels, results in more potential peptides (even after binding affinity filtering). Binding affinities among antigen sources had medians between 100 - 250 nanomolar with the exception of CTA/Self-antigens due to the manual filtering to peptides with binding affinities under 50 nanomolar (Figure 2a). Interestingly, binding affinity medians vary among patients within tumor antigen source and binding affinity variance is especially high in ERV and Virus-derived peptides (Figure 2b). This high variance is likely explained again by the large number of peptides generated from these workflows resulting in more opportunities for binding affinity variability.

**Fig. 2:**
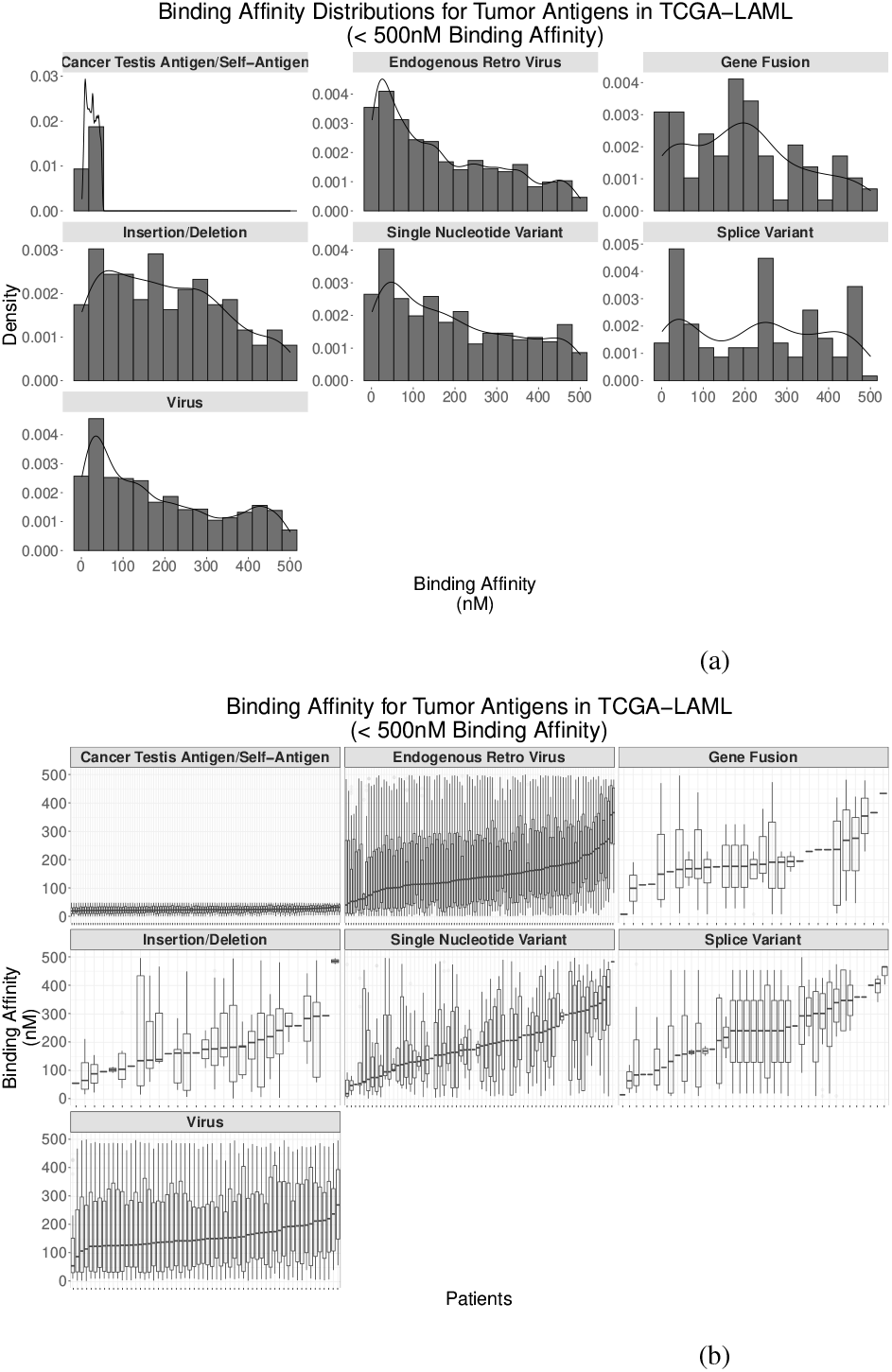
Binding affinities by tumor antigen source among TCGA-LAML patients: All tumor antigen sources showed binding affinity medians between 100 to 250 nanomolar. CTAs/Self-antigens shows markedly lower binding affinity due to the stringent filtering of affinities below 50 nanomolar to compensate for the greater number of peptides generated.

Binding stability distributions within tumor antigen sources also show the expected pattern of an elevated number of low stability pMHC with a long tail of rarer, but more stable pMHCs. Binding stabilities were limited to a maximum of 5 hours for interpretative and illustrative purposes, but some pMHCs, primarily CTA/Self-Antigen-derived and ERV-derived, have estimated binding stability half-lives of up to 66.6 hours. Binding stabilities appear to follow similar trends among patients across tumor antigen sources (Figure 3a). Notably, several splice variant-derived pMHCs show identical distributions of pMHC binding affinity and stability. These patients all appear to have an expressed and detectable retained intron in the *PRTN3* gene.

**Fig. 3:**
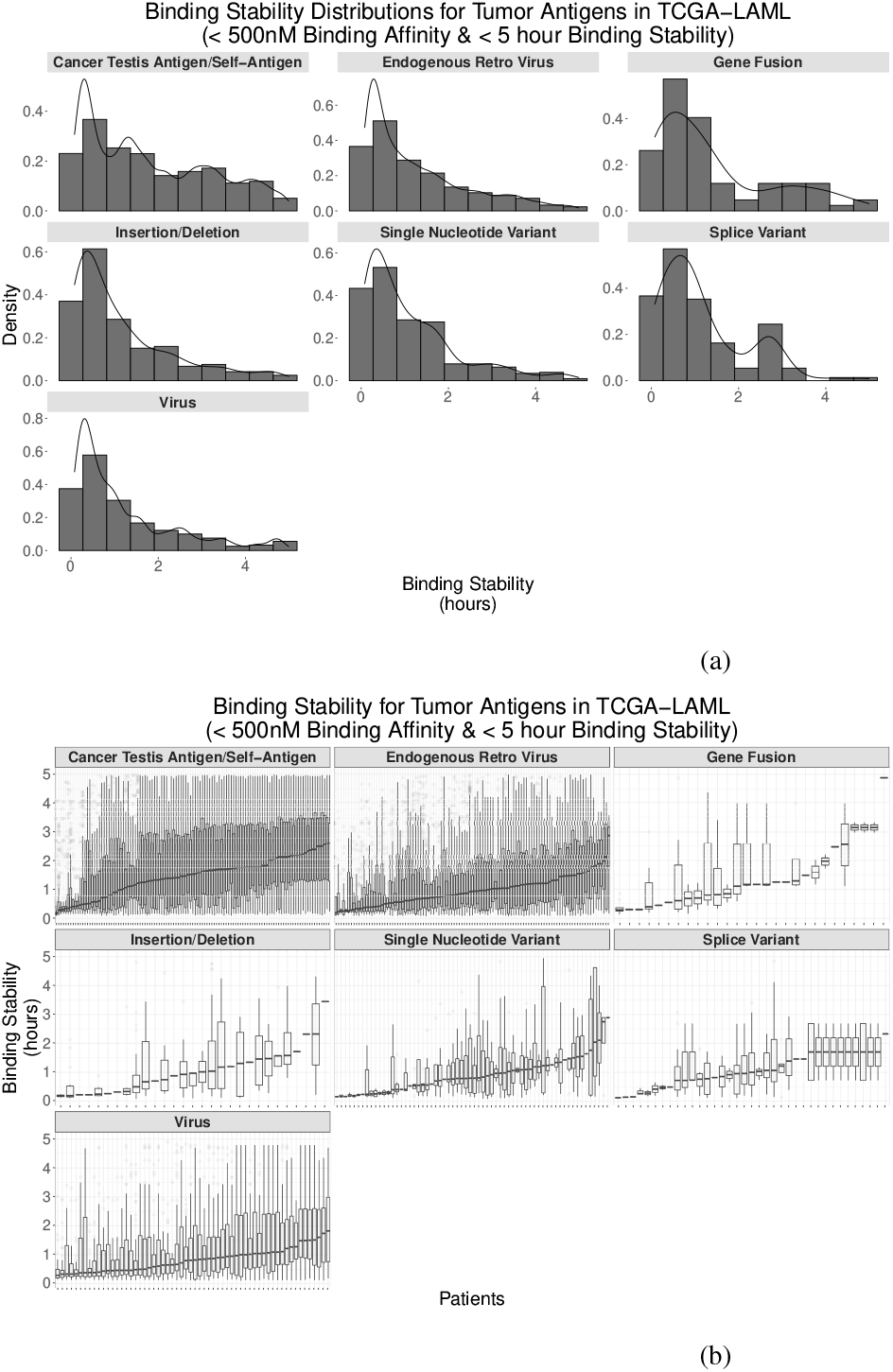
Binding stabilities by tumor antigen source among TCGA-LAML patients: Binding stabilties showed positively skewed distributions with primarily low stability pMHCs and some high stability pMHCs. Note the five hour upper bound used for figure interpretation. This filter excluded a handful of exceptionally high stability pMHCs (up to 66.6 hours).

RNA-based peptide support (calculated as the frequency of peptide coding sequences occurring within tumor-derived RNA sequencing reads) likewise shows variability among antigen sources and among patients (Figures 4). Peptides derived from somatic mutations (e.g. SNVs, InDels, and fusions) show generally lower read support than peptides generated through aberrantly expressed sources (ERVs, CTAs/self-antigens, and splice variants). Both CTA/self-antigens and ERVs show more normal distributions compared to the right-skewed distributions resulting from gene fusions and SNVs. This is likely due to a combination of factors including the averaging effect of having multiple peptides from a single transcript for ERVs and CTA/Self-antigens as well as possible technical artifacts driving SNVs and gene fusions with low read support. Future work into the optimization of each tumor antigen workflow will allow for an improved interpretation of these distributions. More detail on antigen class-specific results are available in the Supplementary Material.

**Fig. 4:**
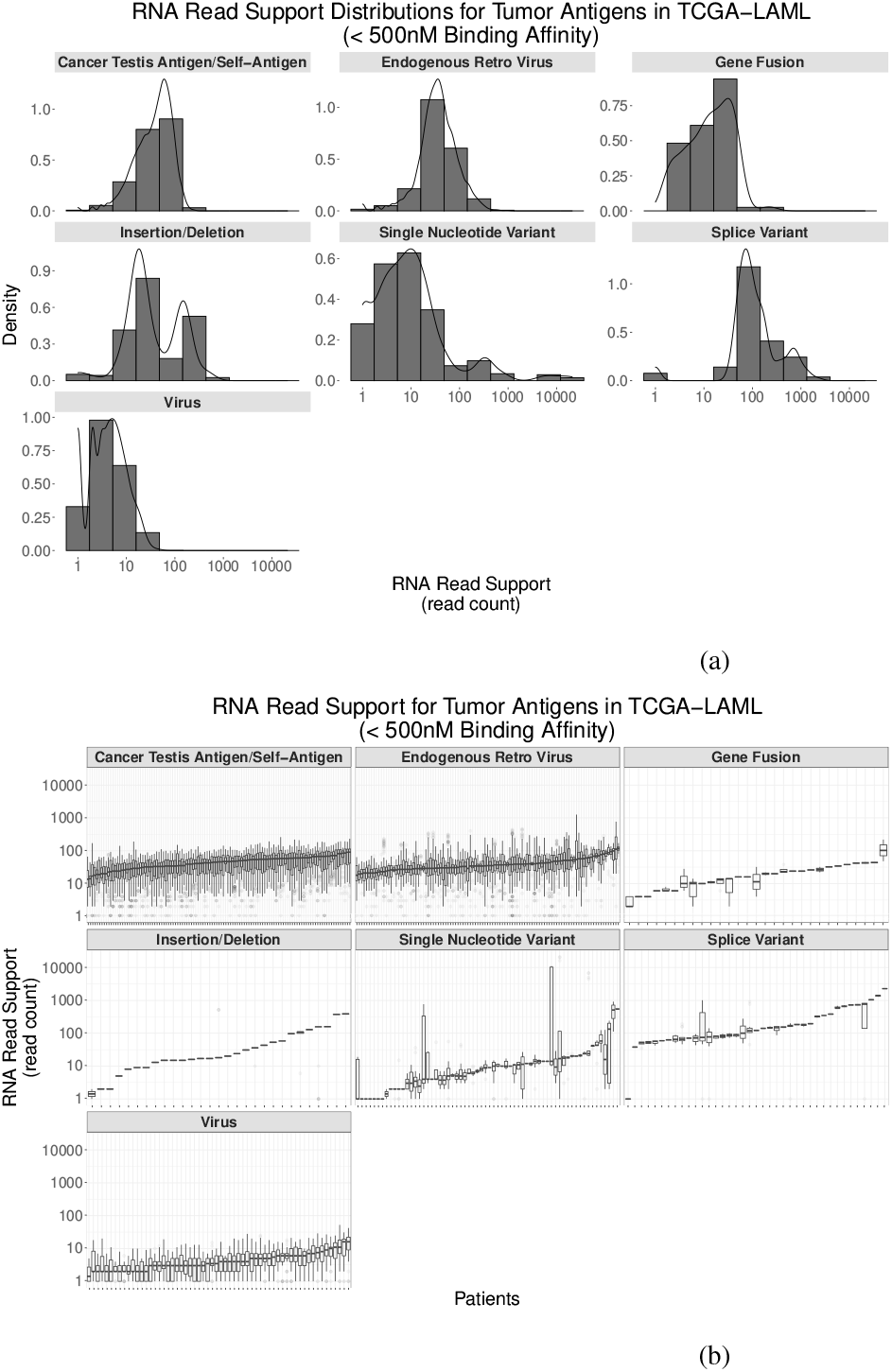
Relative peptide abundance by tumor antigen source among TCGA-LAML patients: Relative peptide abundance (measured by peptide coding sequence detection in RNA sequencing reads) varies across antigen source.

### 3.3 Benchmarking with TESLA

The dataset of experimentally validated immunogenic class I peptides from non-small cell lung cancer (NSCLC) and melanoma patients generated by the TESLA Consortium has allowed for an improved understanding of the factors driving neoantigen immunogenicity (Wells *et al*., 2020). These data also provide opportunities to benchmark the performance of neoantigen workflows (Rieder *et al*., 2022). The set of validated immunogenic peptides was derived from somatic variants, so it does not include the non-somatic variant tumor antigens that LENS predicts. Nonetheless, ensuring a workflow can detect immunogenic peptides serves as a check for correctness of the SNV and InDel neoantigen prediction steps.

LENS was run on the 8 available TESLA patients (2 NSCLC and 6 melanoma) using a union of somatic variant calls from MuTect2, Abra2, Strelka2, and VarScan2. LENS detected 36 of 38 immunogeic peptides from TESLA with 33 peptides passing filtering for reporting in the LENS report. Notably, LENS discovered hundreds of non-SNV/Indel tumor antigens for each patient. More detail can be found in the Supplemental Material.

## 4 Discussion

Predicting the full suite of tumor antigens from genomics data remains a formidable challenge. We developed LENS, coupled with the RAFT framework, to improve upon current workflow offerings in a variety of ways: 1) LENS allows for modularity and extendibility through the Nextflow DSL2 language, 2) LENS explores more tumor antigen sources than previously considered by other workflows, 3) LENS harmonizes peptide abundance estimates among various tumor antigen sources, and 4) LENS includes an application for visualization and interactive exploration to support prioritization of predicted neoantigens for therapeutic applications. LENS is open source, and we expect it to expand through both introduction of new modules, tools, and reference datasets, along with refinement of its tumor antigen prioritization capabilities. There are several objectives discussed below concerning these goals.

A primary objective for the improvement of LENS is expanding the set of supporting sequencing technologies. LENS currently supports data generated through short-read technologies such as the Illumina platform. Illumina’s sequencing-by-synthesis (SBS) chemistry sees widespread usage and support, both within the immuno-oncology community and more broadly throughout the life sciences research community. Despite its popularity, the short read and bulk cell approaches commonly used result in reduced information compared to long read and single cell approaches. We aim to address these limitations within LENS by supporting long read and single-cell sequencing technologies. Long read sequencing technologies allow improved resolution of structural variation that may be relevant to neoantigen prediction. Additionally, single cell sequencing circumvents the confounding effects of bulk sequencing which allows improved understanding of tumor heterogeneity. Empirically deciphering this heterogeneity is crucial to properly prioritizing clonal tumor antigens while also mapping co-occurring subclonal neoantigens to optimize therapeutic targeting.

We plan to further expand LENS through inclusion of third-party reference data and additional bioinformatics tools to provide information about the support (or lack thereof) of a peptide’s immunogenicity. The most important and immediate addition will be the inclusion of Class II binding affinity prediction. We also plan to extract relevant summarized data from large datasets, additional tools, or smaller scale dataset-specific observations in the literature.

Beyond the inclusion of technologies and data, there is also room for improvements within LENS in its current form. LENS supports a variety of tumor antigen sources, but currently it effectively treats these workflows independently. This independence among tumor antigen workflows in LENS also does not allow potentially useful “crosstalk” between workflows. For example, LENS may be able to provide higher confidence for splice antigen targets if there is evidence of somatic splice site variant in the corresponding variant data. While many challenges remain in tumor antigen prediction, we expect that the breadth of features, flexibility, modularity, and ease of use of LENS will support wide adoption as a springboard towards iterative improvements as more data, tools, and an improved understanding of antigen presentation and immunogenicity become available.

## Data Availability

The data underlying this article were accessed from the Genomic Data Commons Data Portal (https://portal.gdc.cancer.gov/). The derived data generated in this research will be shared on reasonable request to the corresponding author.

## References

Almeida, L. G. et al. (2009). Ctdatabase: a knowledge-base of high-throughput and curated data on cancer-testis antigens. Nucleic acids research, 37(suppl_1), D816–D819.

Bjerregaard, A.-M. et al. (2017). Mupexi: prediction of neo-epitopes from tumor sequencing data. Cancer Immunology, Immunotherapy, 66(9), 1123–1130.

Bonaventura, P. et al. (2022). Identification of shared tumor epitopes from endogenous retroviruses inducing high-avidity cytotoxic t cells for cancer immunotherapy. Science Advances, 8(4), eabj3671.

Bright, R. K. et al. (2014). Overexpressed oncogenic tumor-self antigens. Human Vaccines & Immunotherapeutics, 10(11), 3297–3305.

Carreno, B. M. et al. (2015). A dendritic cell vaccine increases the breadth and diversity of melanoma neoantigen-specific t cells. Science, 348(6236), 803–808.

Caushi, J. X. et al. (2021). Transcriptional programs of neoantigen-specific til in anti-pd-1-treated lung cancers. Nature, 596(7870), 126–132.

Chai, S. et al. (2022). NeoSplice: a bioinformatics method for prediction of splice variant neoantigens. Bioinformatics Advances, 2(1).

Cheever, M. A. et al. (2009). The prioritization of cancer antigens: a national cancer institute pilot project for the acceleration of translational research. Clinical Cancer Research: An Official Journal of the American Association for Cancer Research, 15(17), 5323–5337.

Delgado, A. P. et al. (2014). Open reading frames associated with cancer in the dark matter of the human genome. Cancer genomics & proteomics, 11(4), 201–213.

Di Tommaso, P. et al. (2017). Nextflow enables reproducible computational workflows. Nature biotechnology, 35(4), 316–319.

Favero, F. et al. (2015). Sequenza: allele-specific copy number and mutation profiles from tumor sequencing data. Annals of Oncology, 26(1), 64–70.

Filley, A. C. and Dey, M. (2017). Dendritic cell based vaccination strategy: an evolving paradigm. Journal of Neuro-Oncology, 133(2), 223–235.

Frankiw, L. et al. (2019). Alternative mRNA splicing in cancer immunotherapy. Nature Reviews. Immunology, 19(11), 675–687.

Gillis, S. and Roth, A. (2020). Pyclone-vi: scalable inference of clonal population structures using whole genome data. BMC bioinformatics, 21(1), 1–16.

Grandi, N. and Tramontano, E. (2018). Human endogenous retroviruses are ancient acquired elements still shaping innate immune responses. Frontiers in immunology, page 2039.

Haas, B. et al. (2017). Star-fusion: fast and accurate fusion transcript detection from rna-seq. BioRxiv.

Haas, B. J. et al. (2019). Accuracy assessment of fusion transcript detection via read-mapping and de novo fusion transcript assembly-based methods. Genome biology, 20(1), 1–16.

Hu, Z. et al. (2021). Personal neoantigen vaccines induce persistent memory t cell responses and epitope spreading in patients with melanoma. Nature medicine, 27(3), 515–525.

Hundal, J. et al. (2020). pvactools: a computational toolkit to identify and visualize cancer neoantigens. Cancer immunology research, 8(3), 409–420.

Jurtz, V. et al. (2017). Netmhcpan-4.0: improved peptide–mhc class i interaction predictions integrating eluted ligand and peptide binding affinity data. The Journal of Immunology, 199(9), 3360–3368.

Kim, S. et al. (2018). Strelka2: fast and accurate calling of germline and somatic variants. Nature methods, 15(8), 591–594.

Kodysh, J. and Rubinsteyn, A. (2020). Openvax: an open-source computational pipeline for cancer neoantigen prediction. In Bioinformatics for Cancer Immunotherapy, pages 147–160. Springer.

Lang, F. et al. (2022). Identification of neoantigens for individualized therapeutic cancer vaccines. Nature reviews Drug discovery, 21(4), 261–282.

Li, L. et al. (2021). Optimized polyepitope neoantigen dna vaccines elicit neoantigen-specific immune responses in preclinical models and in clinical translation. Genome Medicine, 13(1), 56.

Liepe, J. et al. (2018). Why do proteases mess up with antigen presentation by re-shuffling antigen sequences? Current Opinion in Immunology, 52, 81–86.

Lilleby, W. et al. (2017). Phase i/iia clinical trial of a novel htert peptide vaccine in men with metastatic hormone-naive prostate cancer. Cancer Immunology, Immunotherapy, 66(7), 891–901.

Litchfield, K. et al. (2021). Meta-analysis of tumor-and t cell-intrinsic mechanisms of sensitization to checkpoint inhibition. Cell, 184(3), 596–614.

Lowery, F. J. et al. (2022). Molecular signatures of antitumor neoantigen-reactive t cells from metastatic human cancers. Science, 375(6583), 877–884.

Martin, M. et al. (2016). Whatshap: fast and accurate read-based phasing. BioRxiv, page 085050.

McKenna, A. et al. (2010). The genome analysis toolkit: a mapreduce framework for analyzing next-generation dna sequencing data. Genome research, 20(9), 1297–1303.

Mesri, E. A. et al. (2010). Kaposi’s sarcoma and its associated herpesvirus. Nature Reviews. Cancer, 10(10), 707–719.

Mitchell, G. et al. (2021). Targeting cancer testis antigens in synovial sarcoma. Journal for ImmunoTherapy of Cancer, 9(6), e002072.

Mose, L. E. et al. (2019). Improved indel detection in dna and rna via realignment with abra2. Bioinformatics, 35(17), 2966–2973.

Nakagawa, S. and Takahashi, M. U. (2016). geve: a genome-based endogenous viral element database provides comprehensive viral protein-coding sequences in mammalian genomes. Database, 2016.

Obara, W. et al. (2017). A phase i/ii study of cancer peptide vaccine s-288310 in patients with advanced urothelial carcinoma of the bladder. Annals of Oncology: Official Journal of the European Society for Medical Oncology, 28(4), 798–803.

Ott, P. A. et al. (2017). An immunogenic personal neoantigen vaccine for patients with melanoma. Nature, 547(7662), 217–221.

Perz, J. F. et al. (2006). The contributions of hepatitis B virus and hepatitis C virus infections to cirrhosis and primary liver cancer worldwide. Journal of Hepatology, 45(4), 529–538.

Rasmussen, M. et al. (2016). Pan-specific prediction of peptide–mhc class i complex stability, a correlate of t cell immunogenicity. The Journal of Immunology, 197(4), 1517–1524.

Richman, L. P. et al. (2019). Neoantigen dissimilarity to the self-proteome predicts immunogenicity and response to immune checkpoint blockade. Cell systems, 9(4), 375–382.

Rieder, D. et al. (2022). nextneopi: a comprehensive pipeline for computational neoantigen prediction. Bioinformatics, 38(4), 1131–1132.

Robinson, M. D. et al. (2010). edger: a bioconductor package for differential expression analysis of digital gene expression data. bioinformatics, 26(1), 139–140.

Rolfs, Z. et al. (2018). Global identification of post-translationally spliced peptides with neo-fusion. Journal of proteome research, 18(1), 349–358.

Sahin, U. et al. (2017). Personalized rna mutanome vaccines mobilize poly-specific therapeutic immunity against cancer. Nature, 547(7662), 222–226.

Selitsky, S. R. et al. (2020). Virus expression detection reveals rna-sequencing contamination in tcga. BMC genomics, 21(1), 1–11.

Smith, C. C. et al. (2019a). Alternative tumour-specific antigens. Nature Reviews Cancer, 19(8), 465–478.

Smith, C. C. et al. (2019b). Endogenous retroviral signatures predict immunotherapy response in clear cell renal cell carcinoma. The Journal of clinical investigation, 128(11), 4804–4820.

Smith, C. C. et al. (2019c). Machine-learning prediction of tumor antigen immunogenicity in the selection of therapeutic epitopes. Cancer immunology research, 7(10), 1591–1604.

Tashiro, H. and Brenner, M. K. (2017). Immunotherapy against cancer-related viruses. Cell Research, 27(1), 59–73.

Wang, Y. et al. (2021). Gene fusion neoantigens: Emerging targets for cancer immunotherapy. Cancer Letters, 506, 45–54.

Wells, D. K. et al. (2020). Key parameters of tumor epitope immunogenicity revealed through a consortium approach improve neoantigen prediction. Cell, 183(3), 818–834.

Yang, W. et al. (2019). Immunogenic neoantigens derived from gene fusions stimulate T cell responses. Nature Medicine, 25(5), 767–775.

Zhang, M. et al. (2018). Rna editing derived epitopes function as cancer antigens to elicit immune responses. Nature communications, 9(1), 1–10.

Zhang, Y. et al. (2021). Alternative splicing and cancer: a systematic review. Signal Transduction and Targeted Therapy, 6(1), 1–14.

Zhao, X. et al. (2021). Targeting neoantigens for cancer immunotherapy. Biomarker Research, 9(1), 61.

