## Supplement for "LENS: Landscape of Effective Neoantigens Software"

### LENS
### Lanscape of Effective Neoantigens Software Supplement

**5 Supplemental Tables** **26**

### 1 Supplemental Workflow Figures

#### 1.1 Pre-Processing

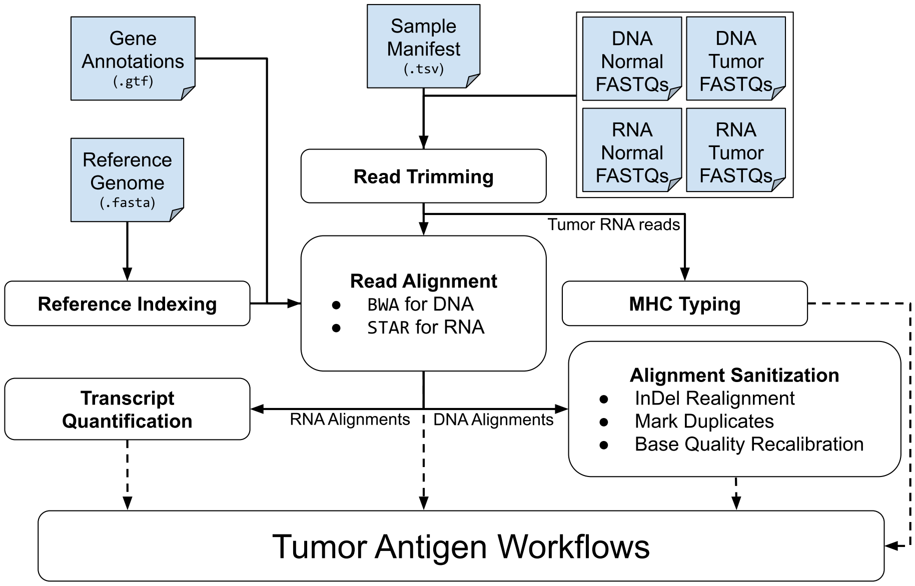

**Supplemental Fig. 1: Pre-processing workflow:** A patient's DNA tumor, RNA tumor, and DNA normal FASTQs are trimmed and aligned to a reference, sanitization steps are performed, and intermediate files are generated for downstream tumor antigen workflows.

#### 1.2 Single Nucleotide Variants and Insertions/Deletions

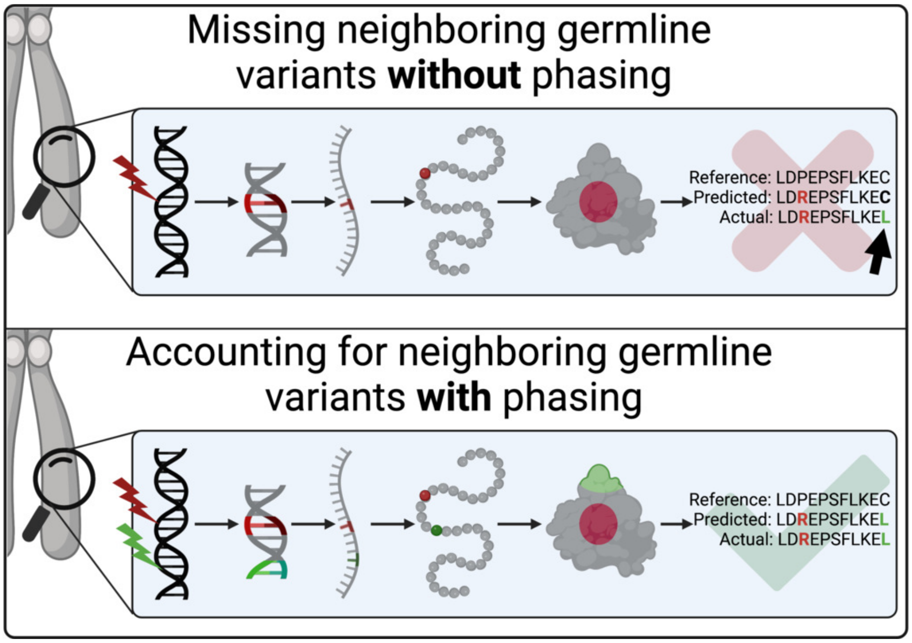

**Supplemental Fig. 2: Combining of somatic variants with neighboring germline variants:** Somatic variants of interest are combined with phased neighboring germline variants during peptide generation. Frameshift InDel variants are combined with phased neighboring germline variants and downstream homozygous germline variants. This strategy maximizes the probability of predicted peptides reflecting those contained within a patient's tumors.

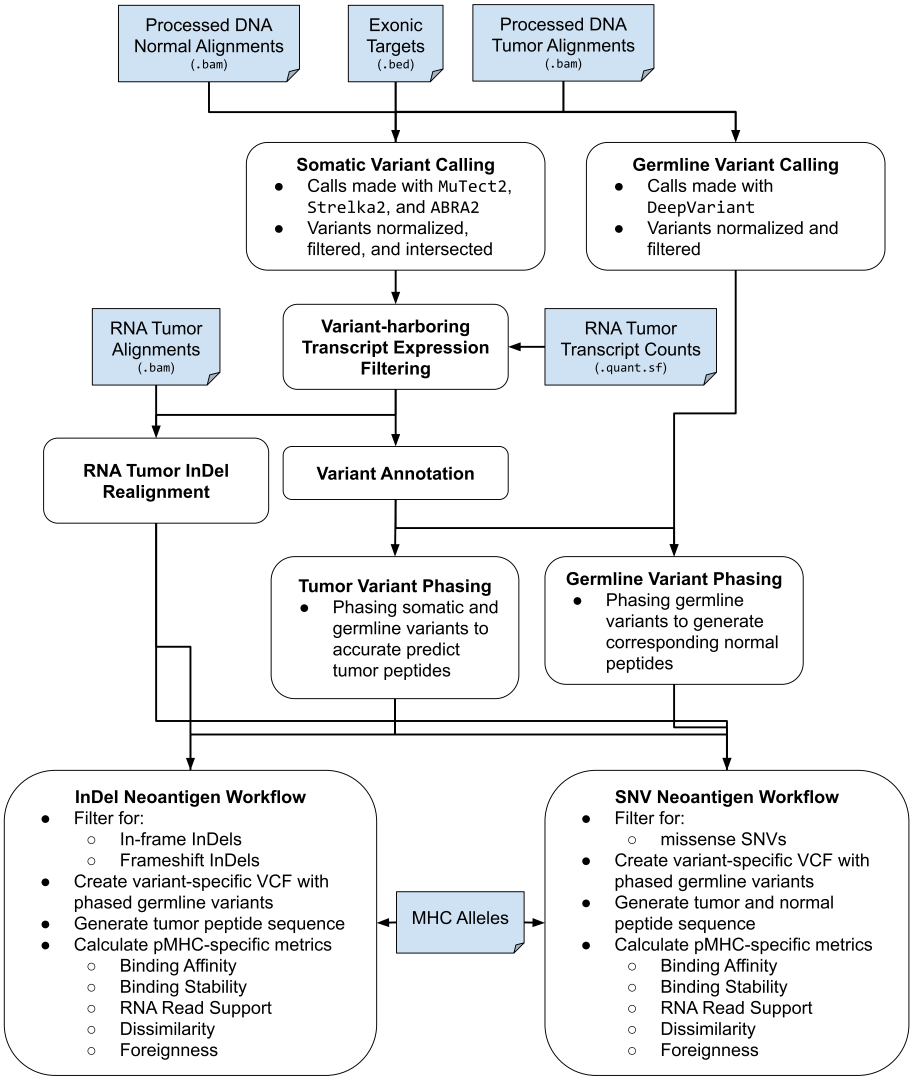

**Supplemental Fig. 3: Single Nucleotide Variant, Insertion, and Deletion workflow:** Somatic variants are called using three separate variant callers: MuTect2, Strelka2, and ABRA2. Germline variants are called using DeepVariant. Somatic variant harboring transcripts are filtered for expression, and the somatic variant and neighboring phased germline variants are incorporated into its coding sequence prior to peptide generation.

#### 1.3 Splice Variants

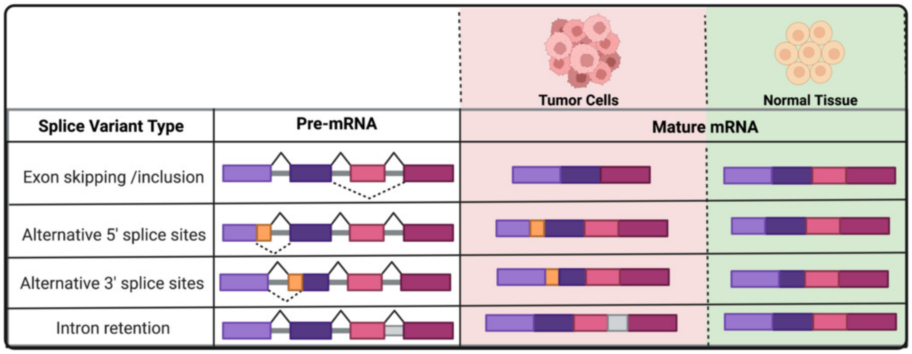

**Supplemental Fig. 4: Examples of tumor splice variants:** Splice variants can present as skipped exons, retained introns, or alternate splice sites. *Adapted from "mRNA Splicing Types", by BioRender.com (2022). Retrieved from https://app.biorender.com/biorender-templates*

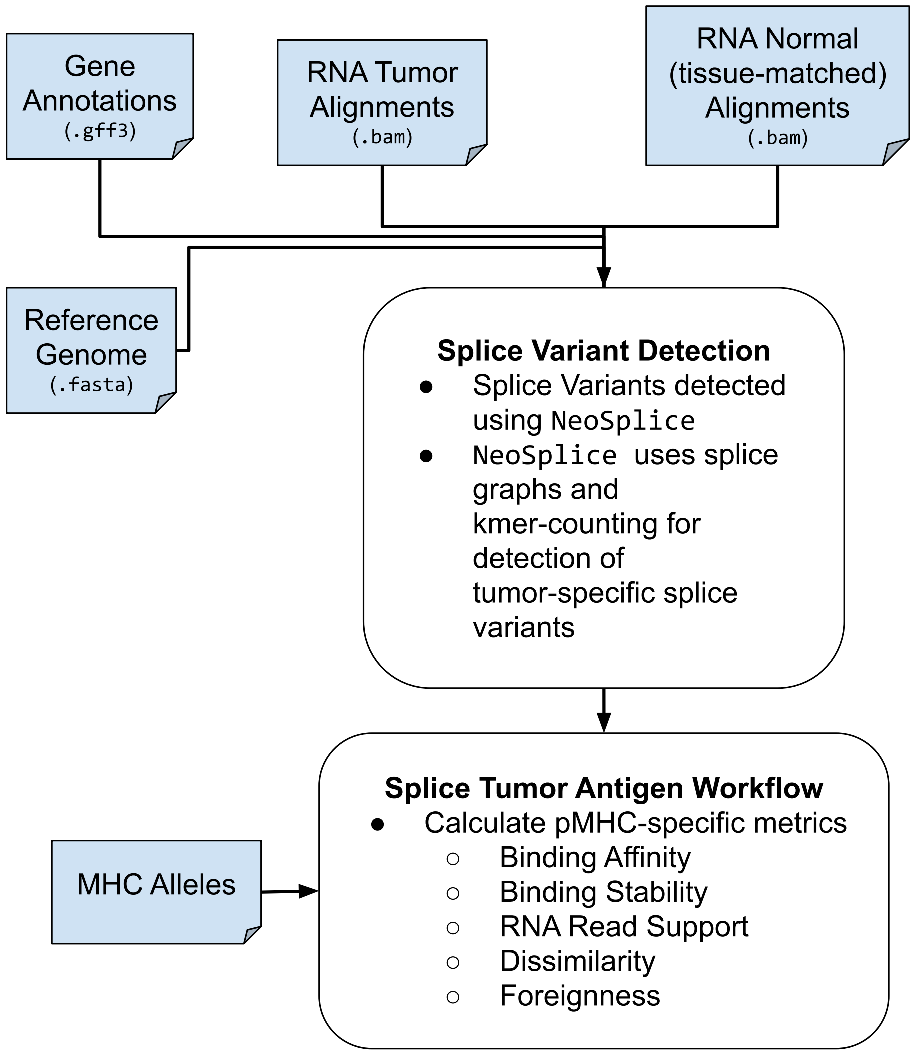

**Supplemental Fig. 5: Splice variant tumor antigen workflow:** The splice workflow relies upon NeoSplice for splice variant detection. NeoSplice considers k-mer distributions between the patient's RNA tumor sample and a tissue-matched control sample.

##### 1.4 Fusion Events

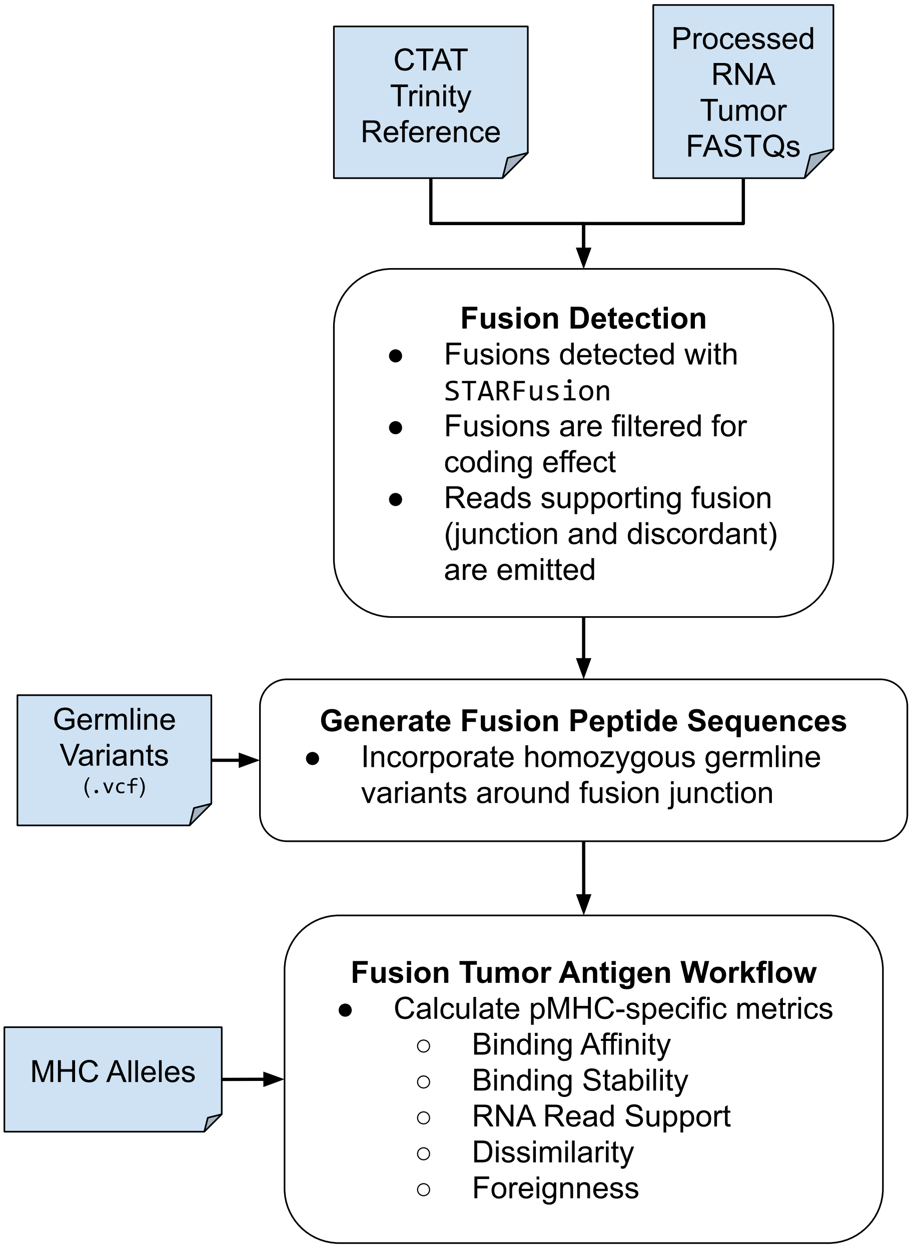

**Supplemental Fig. 6: Fusion variant tumor antigen workflow:** The fusion workflow utilizes STARFusion for fusion detection. Fused transcripts have homozygous germline variants integrated into their sequence prior to translation.

#### 1.5 Tumor-specific Viruses

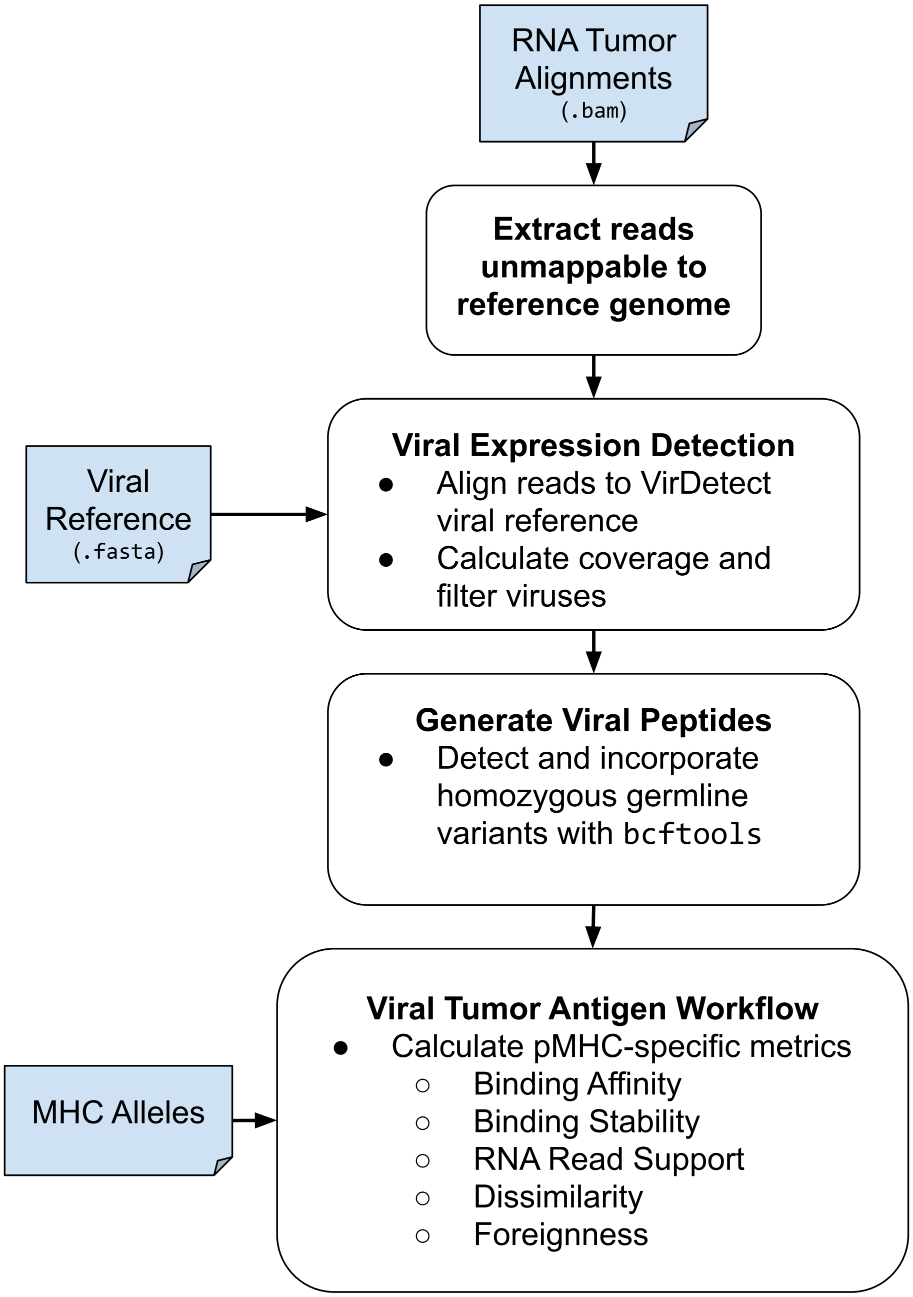

**Supplemental Fig. 7: Viral tumor antigen workflow:** The viral workflow uses the VirDetect reference and alignment strategy to detect viruses expressed within the patient's tumor. Homozygous germline variants are integrated into the viral sequence prior to translation.

#### 1.6 Endogenous Retroviral Elements

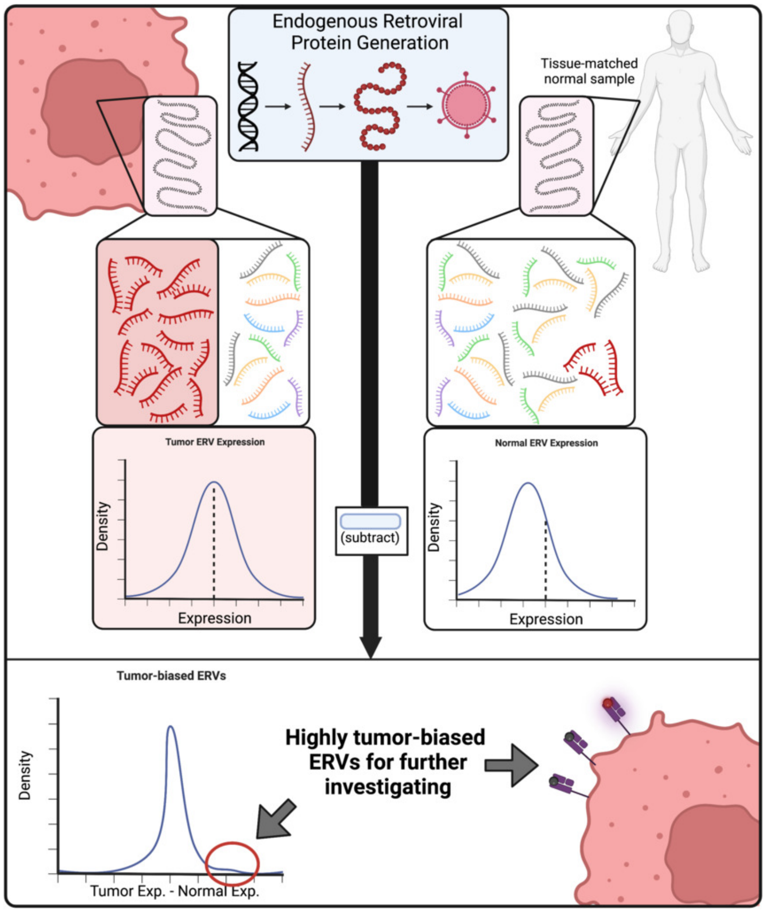

**Supplemental Fig. 8: ERV expression filtering strategy:** ERVs are filtered by comparing the tumoral expression to expression in a tissue-matched normal sample. Only ERVs that show higher expression relative to the normal tissue are considered for processing. *Created with BioRender.com.*

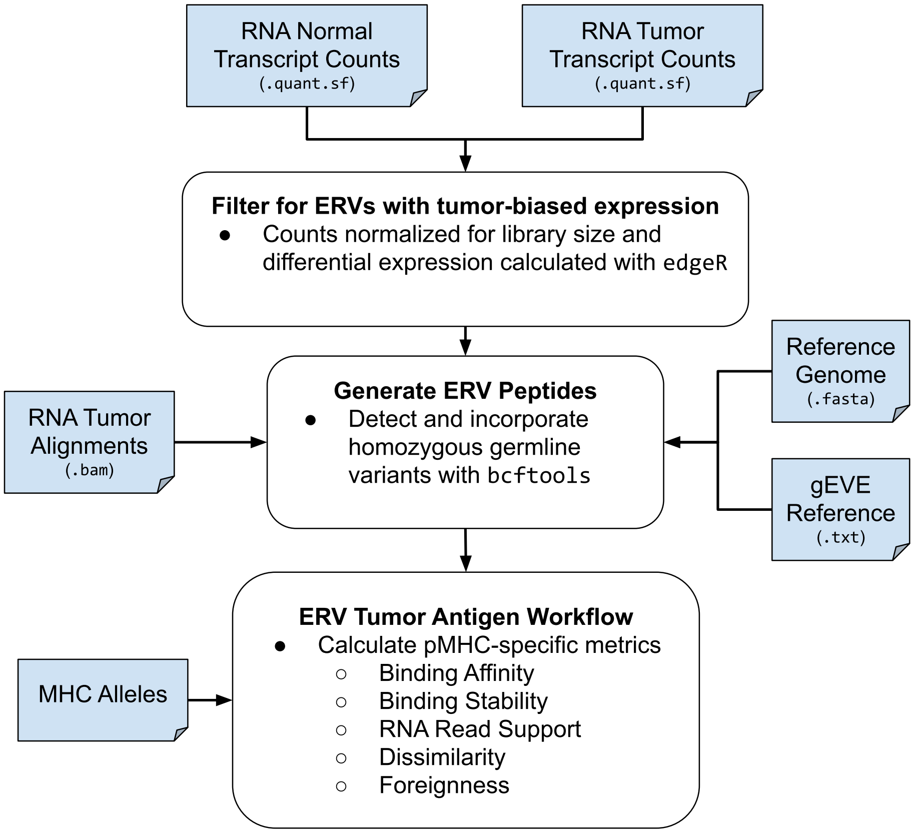

**Supplemental Fig. 9: ERV tumor antigen workflow:** The ERV workflow utilizes predicted ERV ORFs from the gEVE database. Homozygous germline variants are integrated into the coding sequences prior to translation.

#### 1.7 Cancer Testis Antigens and Self-Antigens

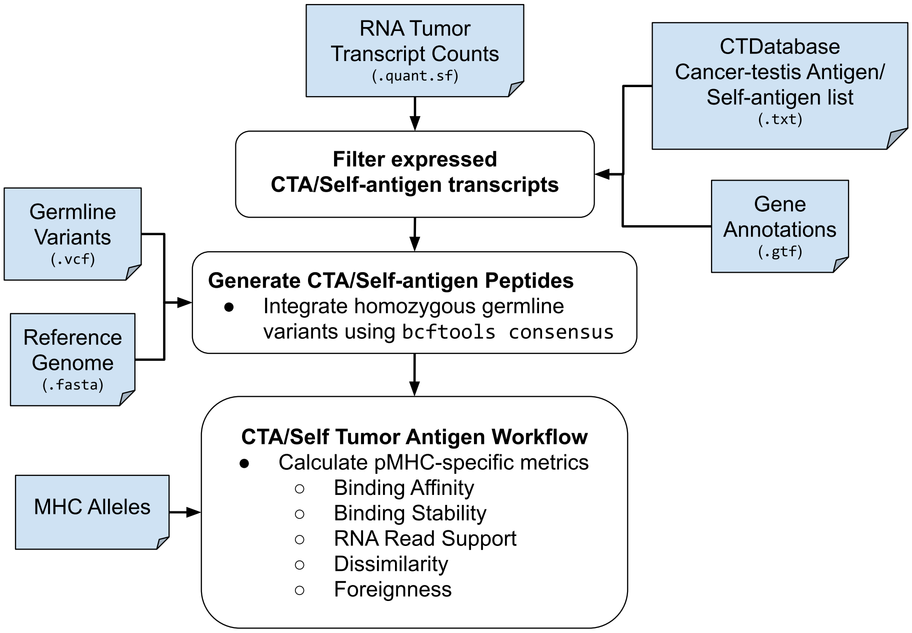

**Supplemental Fig. 10: CTA/Self-antigen tumor antigen workflow:** Cancer testis antigens and self-antigens are processed through the same workflow. Specifically, they are filtered for expression and have homozygous germline variants integrated into their coding sequences prior to further processing.

### 2 Miscellaneous Supplemental Figures

#### 2.1 LENS Viz

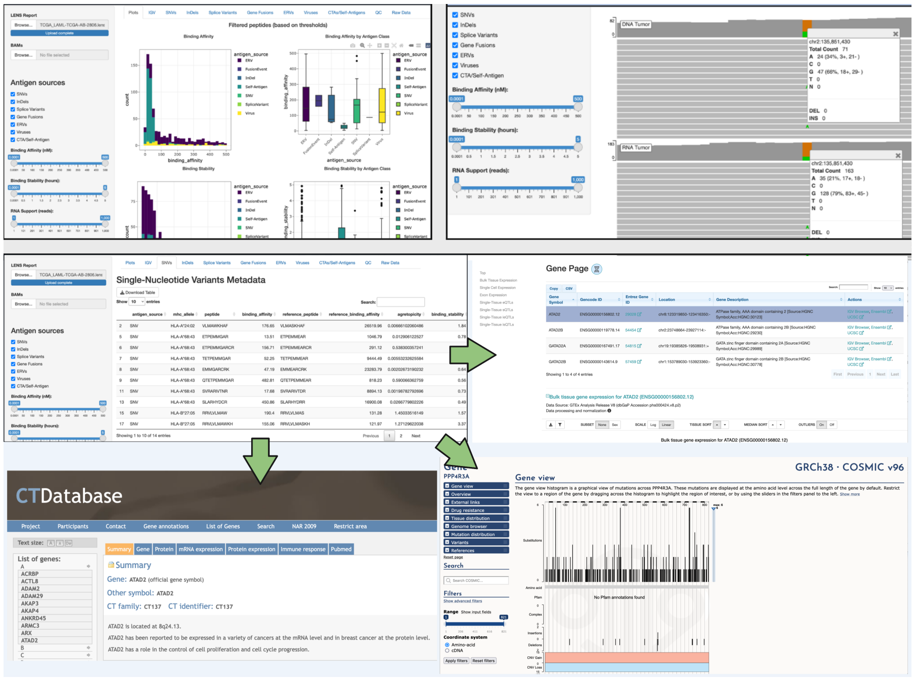

**Supplemental Fig. 11: LENS Viz – A tool for visualizing LENS outputs:** LENS Viz is a shiny app developed to aid users in visualizing and exploring patient-level LENS outputs. LENS Viz allows users to adjust filtering thresholds for various metrics and to interactively view genomic origins of various peptides.

#### 2.2 ERV ORF Filtering

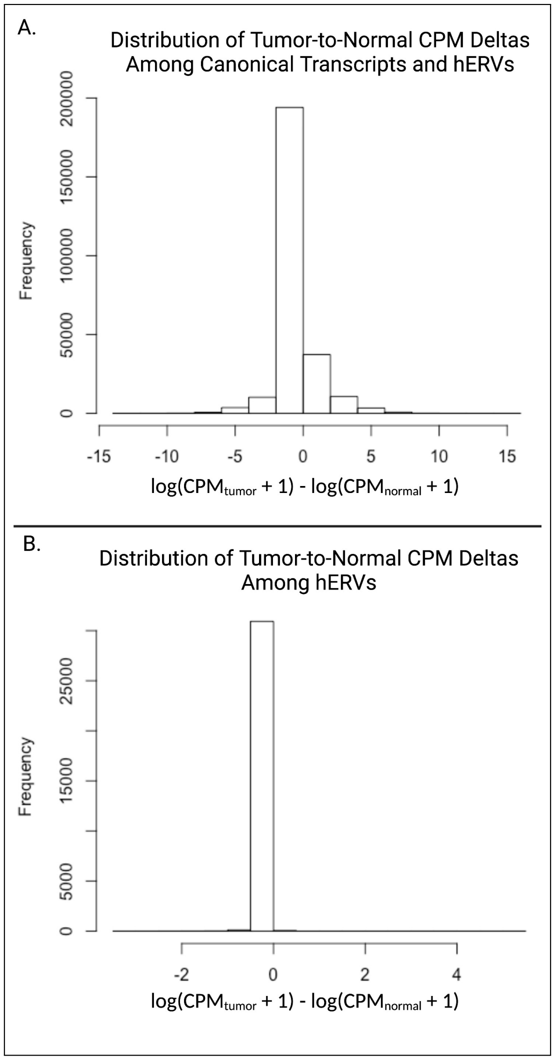

**Supplemental Fig. 12: Endogeous Retrovirus Filtering Strategy:** Recent work has shown endogenous retroviruses can be detectably expressed in normal tissues (She *et al.*, 2022). LENS filters ERVs with relatively high expression in normal tissues (relative to tumors) in an attempt to avoid predicted peptides that may not initiate an immunogenic response due to immune system tolerance. Here we show the EdgeR library-size adjusted CPM values for a melanoma (tumor) and skin (normal) sample. Subfigure **A** contains the distribution of tumor-to-normal expression differences among all canonical transcripts and hERVs. Subfigure **B** shows the distribution of tumor-to-normal expression differences among all hERVs. LENS utilizes a relatively stringent filter of $log(CPM_{tumor} + 1) - log(CPM_{normal} + 1) > 1$ to optimize for ERVs with tumor-biased expression.

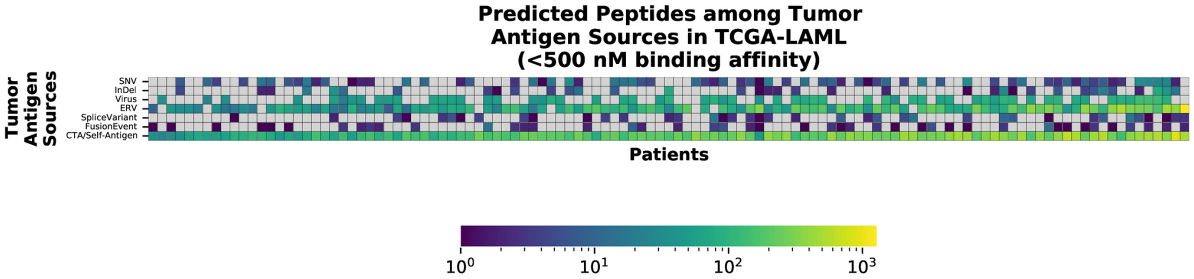

**Supplemental Fig. 13: TCGA-LAML tumor antigen distribution by patient:** Tumor antigen distributions across 115 TCGA-LAML patients. Predicted neoantigens range between 17 to 1,153 peptides with a median of 177 peptides per patient.

### 3 Supplemental Results

#### 3.1 SNV and InDel Neoantigens in TCGA-LAML

Despite low tumor mutational burden in AML, some SNV neoantigens have been previously identified. Greif et al. reported five tumor-specific non-synonymous mutations in a single AML sample via whole transcriptome sequencing (Greif *et al.*, 2011). InDels have also been identified within AML. For example, Lee et al. reported a novel InDel within the *KIT* gene, a commonly mutated gene in AML patients, in a clinical sample from a 35 year old female (Lee *et al.*, 2016). Two neoantigens derived from *NPM1* mutations have been identified (van der Lee *et al.*, 2019, 2021). Variant calling was performed on TCGA-LAML samples using a consensus approach in which each variant had to be detected by at least two separate variant callers (for SNVs) or three separate variant callers (InDels) to be considered. Furthermore, variants had to exceed relative transcript abundance upper quartile and the nucleotide sequence surrounding the SNV or InDel variant of interest must also have been discoverable within the patient's corresponding RNA sequencing reads. As a result, a patient's SNV and InDel neoantigen peptide count is expected to the proportional to, but less than, the total number of high confidence somatic variant calls observed within the DNA data.

We discovered potentially targetable somatic SNVs in 95 of 115 patients which translated into SNV neoantigen targets for 64 patients with a median of 5 SNV-derived neoantigen peptides (min: 1, max: 31). Several genes were observed to contain targetable SNVs in multiple patients including *DNMT3A* in eight patients, *FLT3* in four patients, *TP53* in three patients, as well as *RUNX1* and *PTPN11* in two patients. These genes have been previously observed to have relatively high mutation rates in AML cancers. Most genes with targetable SNVs (108/120, 90%) were private to single patients. One SNV-derived peptide, **YTDVSNMSCL**, from *DNMT3A* was observed within two patients. No other SNV-derived tumor antigens were shared among patients; however, this observation is not surprising given the previously observed low somatic mutation rate within AML patients (Kandoth *et al.*, 2013).

InDel-derived neoantigens showed a similar pattern among patients with 61 of 115 patients having potentially targetable InDels. Thirty-one of the 63 patients had InDel-derived peptides predicted with a median of 3 peptides per patient. Interestingly, two patients are outliers with 52 and 56 InDel-derived peptides each. These peptides were generated by frameshifts in 100% and 91.1% of cases, respectively. This provides an example of the potency of frameshift mutations in the context of peptide generation. Several InDel-harboring loci were shared among individuals including *RUNX1*, *NPM1*, and *FLT3* which were all found in 3 sets of patients. Two peptides, **AIQDLCLAV** and **QEAIQDLCL**, were discovered in 3 patients and two patients, respectively. Both peptides are generated through a common frameshift mutation in *NPM1*, W288Xfs*12. No other peptides were shared among patients.

#### 3.2 Splice Variant Neoantigens in TCGA-LAML

Given the relatively low mutational burden observed within AML samples, other potential tumor antigen sources are of great interest. Splice variants associated with mutations in splicing-associated proteins have previously been identified for AML, including possible driver mutations in genes such as *L-Myc* and *PTPN6* (Zhou and Chng, 2017). We tested patients for tumor-specific splice variants using NeoSplice coupled with a non-matched normal blood sample(Chai *et al.*, 2022a).

Our NeoSplice analysis discovered splice-derived tumor antigens in 39 of the 115 patients with a median splice tumor antigen count of 3 (min: 1, max: 16). Several loci harboring tumor splice antigens tended to be shared among patients, including *STAB1* (14 patients) and *PRTN3* (14 patients), among others. Interestingly, the patients with the *PRTN3* splice variant all retained the intron between exons 3 and 4. The resulting intronic sequence coded for three peptides shared by most of the 14 patients, **ILLIQLSSPA**, **KLNDILLIQL**, and **KLNDILLIQLS**. Previous work shows one of the splice antigen-generating loci, *Myeloperoxidase* (*MPO*), can have a splice site substitution variant that results in the retention of intron 11 in some AML patients (Li *et al.*, 2019).

Some genes, like *SLCA11A1* and *NCF1*, were observed to have multiple tumor-associated splice variant peptides within at least one patient, but their cellular functions, however, make it difficult to disentangle between potential tumor expression and evidence of an immune response (Cellier, 2013; Masson *et al.*, 1986).

#### 3.3 Fusion Neoantigens in TCGA-LAML

The ability of fusion genes to create new, tumor-specific transcripts coupled with the potential for frameshifting effects on the protein product make them appealing targets for neoantigen predictions. A recent large-scale study of in-frame fusion events across 539 AML cases documented 296 fusion genes of which

57 were shown to be recurrent. Chen et al.'s results focused on in-frame fusion events, but highlight two important factors for fusion-derived neoantigens: 1) fusion events are not rare within AML cases (82.2% of all acute leukemia cases considered had at least one high-confidence fusion events) and 2) recurrent fusion genes suggest the potential for vaccination of shared fusion events (Chen *et al.*, 2021). LENS can help with understanding the relationship between shared variants among patients and which neoantigens are worth pursuing.

Our TCGA-LAML fusion-derived neoantigen workflow identified a gene fusion event observed in fourteen patients, *KANSL1-ARL17A/B*. This gene fusion has been previously observed in AML patients and is believed to be a predisposition fusion gene; however, this event has been observed in normal tissues as well which suggests it may be a subpar vaccination target (Zhou *et al.*, 2017; López-Nieva *et al.*, 2019). The next most common fusion gene (*RARA-PML*) was observed in five patients. Two fusion genes were observed among three patients each (*RUNX1-RUNX1T1* and *CBFB-MYH11*). There were several shared tumor antigens observed among patients. These include **RANKVSVWR** (9 patients), **IRANKVSVWR** (7 patients), and **QIRANKVSVWR** (7 patients) from a shared *KANSL1-ARL17* gene fusion event. A shared *RUNX1-RUNX1T1* fusion also resulted in peptides **GPREPRSA** and **GPREPRSARA** being observed in two patients.

#### 3.4 Viral and Endogenous Retroviral Tumor Antigens in TCGA-LAML

Oncogenic viruses that are highly expressed within a tumor may serve as a suitable target for tumor vaccination. AML is not canonically associated with viral oncogenesis, however viral and endogenous retroviral elements remain potential sources of tumor antigens in AML. Clinical studies investigating the relationship between hERVs and AML are ongoing (NCT04406207) and recent work using single-cell RNA sequencing discovered a variety of viral and retroviral elements expressed in AML tumors (NCT, 2022).

We discovered some level of retroviral expression in 107 patients. Of these, 56 patients expressed ORFs associated with single ERV-associated proteins, 18 patients expressed ORFs associated with two ERV-associated proteins, and 11 expressed ORFs from three or more ERV-associated proteins. The most abundant hERV type observed among patients was hERV-K10 followed by hERV-E. Several of the most commonly shared peptides are derived from an ERV ORF located on the positive strand of chromosome one from positions 206,057,742 to 206,058,290. These peptides include **AIYQNRLAL** (40 patients), **FPAIGGFKTL** (39 patients), **TLIIRVIIV** (38 patients), and **CLIPVFLQM** (30 patients). Notably, all but one viral peptide were generated from a hERV-K113 sequence contained within the VirDetect viral reference. It's also worth noting the ERV ORFs from the gEVE database are computationally predicted. An orthogonal strategy like Ribosome sequencing may aid in filtering out false-positive ORFs (Nakagawa and Takahashi, 2016).

Both the current ERV and viral workflows produce a multitude of predicted tumor antigen peptides due to the entirety of the viral or retroviral sequence being included. Further work is required to more rigorously prioritize and validate both the expression of these elements and their potential contribution to tumor antigen vaccine strategies.

#### 3.5 Cancer Testis Antigens and Self-antigens in TCGA-LAML

Multiple CTAs have been previously reported for AML, including *Cyclin A1*, *MAGE*, *PASD1*, *PRAME*, and *RAGE-1* (Anguille *et al.*, 2012). The Cancer Testis Antigen Database (http://www.cta.lncc.br/) provides a variety of well-documented testis-specific or testis-elevated expressed transcripts that may serve as suitable vaccination targets within tumors expressing them. We considered the relative expression value of all transcripts associated with genes in the Cancer Testes Database and required CTA transcripts to be relatively highly expressed (above 95th percentile) in a patient's tumor RNA data. Several CTA loci were highly expressed in a number of patients. This includes 115 patients expressing *KIAA0100*, a CTA expressed in hematological cancers and an association with an immune response in AML (Chen *et al.*, 2005). Another potential target, *DCAF12*, was observed to be expressed in 112 patients. *DCAF12* is associated with immune response in some tumor types, but is also involved in erythoid differentiation which suggests it may be a very poor target in AML (An *et al.*, 2014). This highlights the importance of considering tumor context when evaluating antigen sources. Two other intriguing targets, *PRAME* and *SPAG6*, were discovered in 23 patients and 7 patients respectively. *PRAME* expression has been shown to have an immunogenic response in AML (Quintarelli *et al.*, 2011) while *SPAG6* has been observed in AML previously (Steinbach *et al.*, 2006). Notably, CTAs and self-antigen peptides can be generated by the entirety of each transcript's coding sequence. As a result, patients typically have several magnitudes more CTA peptides than other types and will require more stringent filtering prior to prioritization.

The above rudimentary analysis emphasizes the importance of the consideration of a wide breadth of tumor antigen source when prioritizing antigens for vaccination. AML is an extreme case due to its relatively low TMB, but other tumor types would still require inclusion of tumor antigen sources beyond the standard SNV, InDel, and Fusion antigens to comprehensively map the expressed antigen landscape.

#### 3.6 TESLA Benchmarking

The set of experimentally validated immunogenic peptides from the TESLA consortium allows for benchmarking of SNV and InDel derived tumor antigen workflows. We ran LENS on the 8 available TESLA patients (2 NSCLC, 6 melanoma) to validate and benchmark our SNV and InDel subworkflows. LENS provides users with the flexibility of using a variety of variant callers and either a union or intersection strategy. We benchmarked using a union strategy with MuTect2, Strelka2, VarScan2, and Abra2 variant calls. This approach, coupled with relatively low TPM filtering (1st percentile), detected 36 of 38 validated immunogenic peptides and 34 peptides were present in the final LENS output (see Supplemental Table 1). LENS also predicted non-SNV/InDel tumor antigens for each patient – specifically, an average of 2,264, 168, 6, 3, and 177 peptides from CTA/Self-antigens, endogenous retroviruses, gene fusion events, splice variants, and viruses, respectively, among the eight patients. The count for each antigen source for each patient can be found in Supplemental Table 2.

| Patient ID | Allele | Peptide | Variant Type | Detectable | In LENS Report | Variant Coordinates | Comments |
|---|---|---|---|---|---|---|---|
| 1 | A*02:01 | ALDHMFMYFL | SNV | True | True | chr19:1979196 | N/A |
| 1 | A*02:01 | FLDPDLTNI | SNV | True | False | chr17:63479089 | See below[n] |
| 1 | A*02:01 | FLGSLLILV | SNV | True | True | chr18:49258444 | N/A |
| 1 | A*02:01 | FLNCDIMLGV | SNV | True | True | chrY:14830012 | N/A |
| 1 | A*02:01 | KAWENFPNV | SNV | True | False | chr1:171574958 | See below[†] |
| 1 | A*02:01 | RVYDALNLL | SNV | True | True | chr13:113633938 | N/A |
| 1 | A*02:01 | YLYHRVDVI | SNV | True | True | chr16:8899702 | N/A |
| 1 | A*68:01 | DTIDVSKLNR | SNV | True | True | chr3:69071558 | N/A |
| 1 | A*68:01 | EIIPQCIAR | SNV | True | True | chr1:84490400 | N/A |
| 2 | A*01:01 | ILDTAGHEEY | SNV | True | True | chr1:114713907 | N/A |
| 2 | A*02:01 | ALPPTVYEV | SNV | True | True | chrX:154535207 | N/A |
| 2 | A*02:01 | GLYGNLIVL | SNV | True | True | chr1:229539581 | N/A |
| 2 | B*57:01 | KSFKEIKLW | SNV | True | True | chr3:52586598 | N/A |
| 3 | A*01:01 | FTNESYLELY | SNV | True | True | chr1:11080888 | N/A |
| 3 | A*01:01 | LSDGPSMGRY | SNV | True | True | chr4:30724342 | N/A |
| 3 | A*03:01 | AINRPTVLK | SNV | True | True | chr4:122743639 | N/A |
| 3 | A*03:01 | ATRYNYTSEK | SNV | True | True | chr5:157315074 | N/A |
| 3 | A*03:01 | ATYKGVPYEVK | SNV | True | True | chr4:106347631 | N/A |
| 3 | A*03:01 | HLFDIQGLPK | SNV | True | True | chr17:44189398 | N/A |
| 3 | A*03:01 | ILFRTPSVAKV | Unknown | False | False | Unknown | N/A |
| 3 | A*03:01 | KIVEMSTSK | Unknown | False | False | Unknown | See below[o] |
| 3 | A*03:01 | KIYTGEKPYK | SNV | True | True | chr7:64978592 | N/A |
| 3 | A*03:01 | KLRDEISLAK | SNV | True | True | chr11:4074582 | N/A |
| 3 | A*03:01 | RLFPYALHK | SNV | True | True | chr7:131327945 | N/A |
| 3 | A*03:01 | RTREFTAKK | SNV | True | True | chr1:192579165 | N/A |
| 3 | B*08:01 | HALRRHYHL | SNV | True | True | chr10:9433786 | N/A |
| 12 | A*02:01 | YQANVVWKV | INDEL | True | True | chr12:62381609 | N/A |
| 12 | A*03:01 | ALYFNSQWK | SNV | True | True | chr7:101131955 | N/A |
| 12 | B*07:02 | RGRMQTASL | SNV | True | True | chr1:147654318 | N/A |
| 12 | C*06:02 | VRINTARPV | SNV | True | True | chr16:24563640 | N/A |
| 16 | A*02:01 | KLLSFHSV | SNV | True | True | chr19:37388827 | N/A |
| 16 | A*02:01 | YLNEAVFNFV | SNV | True | False | chr11:100959900 | See below[†] |
| 16 | B*27:05 | RRSMLFARH | SNV | True | True | chr21:45277127 | N/A |
| 16 | C*05:01 | KTDTGVHATL | SNV | True | True | chr2:189039401 | N/A |
| 9 | A*01:01 | ATDTNNLNVNY | SNV | True | True | chr8:23447838 | N/A |
| 8 | A*01:01 | ISDSLLWKY | SNV | True | True | chr17:5462441 | N/A |
| 9 | A*01:01 | YLDGKVVDY | SNV | True | True | chr5:95520755 | N/A |
| 4 | A*02:01 | ALDPHSGHFV | SNV | True | True | chr12:57751647 | N/A |

**Supplemental Table 1: Validated Immunogenic TESLA peptides detected through intersection filtering method** LENS detected 36 of 38 validated peptides (35 SNVs, 1 INDEL) in 8 TESLA patients using the intersection strategy for somatic variant filtering. Thirty-three peptides were available in the filtered LENS output. The exceptions are described below.
[†]Variant not translated by default due to snpEff warning "WARNING_TRANSCRIPT_NO_START_CODON"
[o]Missense variant results in self peptide as KIVEMSTSK is present in at least 13 canonical protein sequences from EIF5A and EIF5A2.
[n] Causal somatic variant detected, but peptide had no read support from tumor RNA sample.

| Patient ID | CTA/Self-antigen | ERV | Fusion | Splice | Virus |
|---|---|---|---|---|---|
| 1 | 1,824 | 851 | 7 | 12 | 267 |
| 2 | 620 | 183 | 0 | 7 | 0 |
| 3 | 702 | 307 | 1 | 0 | 104 |
| 4 | 2,892 | 0 | 0 | 3 | 244 |
| 8 | 362 | 0 | 0 | 0 | 0 |
| 9 | 67 | 2 | 0 | 0 | 7 |
| 12 | 6,499 | 0 | 26 | 0 | 790 |
| 16 | 5,143 | 0 | 12 | 0 | 0 |

**Supplemental Table 2: Non-SNV/INDEL Tumor Antigens in TESLA Patients** LENS discovered an average of 2,264, 168, 6, 3, and 177 peptides from CTA/Self-antigens, endogenous retroviruses, gene fusion events, splice variants, and viruses, respectively, among the TESLA patients.

### 4 Supplemental Methods

#### 4.1 Harmonized Antigen Quantification Method

Proper quantification of potential tumor antigens is crucial for selecting suitable therapeutic targets. Many of the tools useful for detecting potentially tumor antigen-generating genomic aberrations provide a variety of quantification metrics. For example, Arriba and STAR-Fusion are both used to discover and quantify gene fusion events, but provide different metrics (predicted chimeric read count and fusion fragments per million (FFPM), respectively). Furthermore, neither of these metrics are easily translatable into other common quantification metrics, like transcript per million (TPM) provided by tools like Salmon.

We constructed a harmonized antigen quantification approach in an attempt to provide a metric that allows an understanding of relative abundance of tumor antigens among antigen sources. Methods are largely similar among antigen sources though some source-specific considerations must be accommodated. The resulting metric is effectively a raw read count of reads supporting the tumor antigen.

##### 4.1.1 SNVs and InDels

Both the SNV and InDel workflows share a strategy in which the genomic origin of the targetable peptide-generating somatic variant is identified and reads supporting the variant (and its associated genomic context) are counted from the RNA tumor BAMs. Specifically,

1. SNV-derived and InDel-derived peptides (and their corresponding coding sequences) are generated by LENSTools.

2. The variant's position relative to the start of the coding sequence of the transcript harboring it is used to isolate genomic origin.

3. Reads that overlap the variant position of interest *and* are capable of containing the entirety of coding sequence of the peptide are fetched using PySam (Gilman *et al.* (2019)).

4. Reads from Step 2 are filtered to a set containing the predicted peptide's coding sequence and counted. The resulting sum is provided as reads_with_peptide in the LENS report.

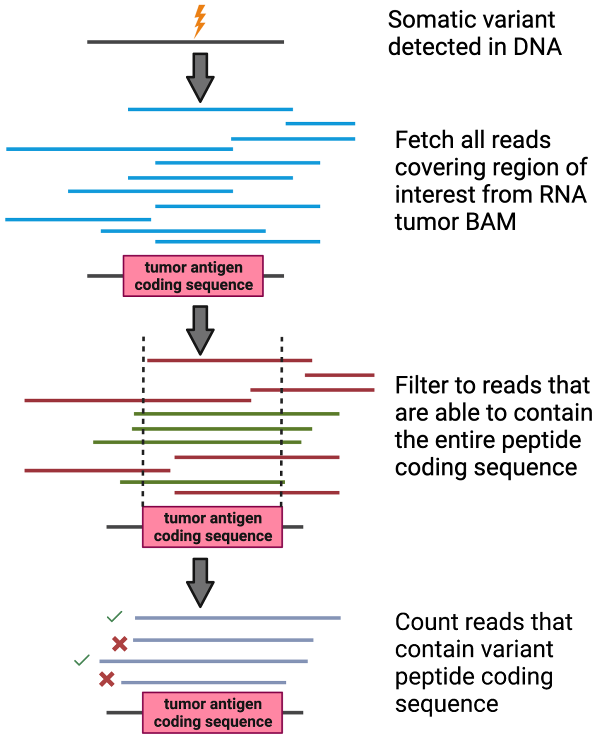

###### 4.1.2 Splice Variants

Splice variant quantification relies upon peptide, coding sequence, and region of interest data from NeoSplice. Specifically,

1. Splice variant peptides, associated coding sequence, and splice region of interest are extracted from NeoSplice outputs.

2. Reads that overlap the splice variant region of interest are fetched from the RNA tumor samples using PySam.

3. Reads are filtered to a reduced set containing the splice variant peptide's coding sequence and counted. The resulting sum is provided as reads_with_peptide in the LENS report.

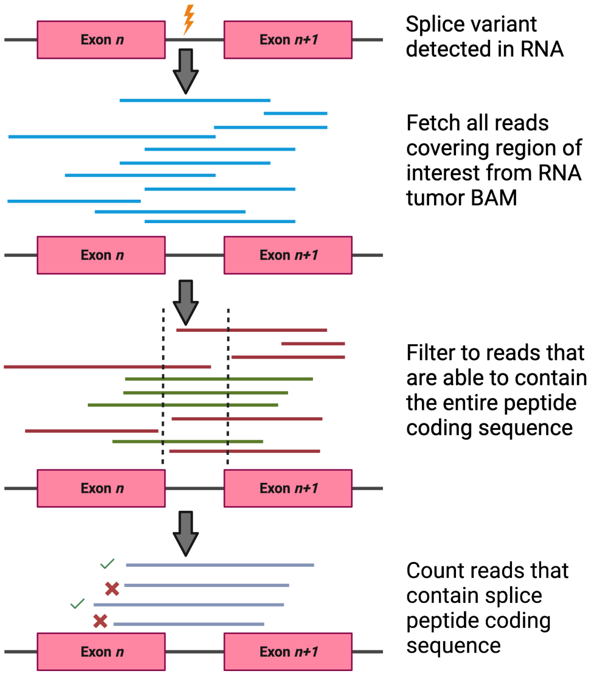

##### 4.1.3 Gene Fusions

The gene fusion quantification method is the most divergent from other tumor antigen sources. Reads derived from a fused transcript product, but not either overlapping the fusion junction point or paired with a discordantly mapped partner cannot be distinguished from reads generated from canonical transcript products. The inability to disentangle these conflated reads results in our fusion peptide estimate being an underestimate of abundance relative to other antigen sources. Specifically,

1. Fusion-derived peptides with favorable metrics (e.g. binding affinity $\leq 1000nM$) have relevant metadata from fusion caller output files.
2. Junction spanning and discordantly mapped reads provided by fusion caller are extracted from tumor RNA FASTQs.
3. Reads are filtered to set that contain peptide coding sequence and are counted. The resulting sum is provided as reads_with_peptide in the LENS report.

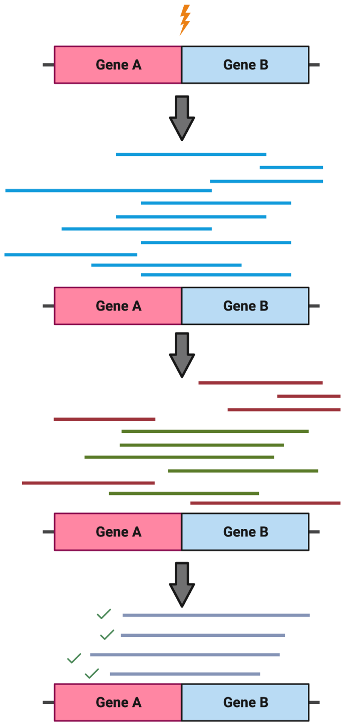

###### 4.1.4 Endogenous Retroviruses, Cancer Testis Antigens, and Self-antigens

Endogenous retroviruses (ERVs), cancer testis antigens (CTAs) and self-antigens are a unique class of tumor antigen sources as their targetable peptides come from aberrant expression rather than byproducts of genomic abnormalities. As a result, the entirety of the translated protein is considered for tumor antigen peptide generation though only the peptide-specific coding sequence is quantified to allow for suitable comparison to other tumor antigen quantification strategies. Specifically,

1. The entirety of an expressed ERV, CTA, or self-antigen protein is processed through NetMHCpan and other pMHC characterization tools.

2. Peptides of interest (e.g. high binding affinity) have their genomic origin identified either through canonical transcript annotation (CTA/self-antigen) or through gEVE database coordinates (ERV).

3. Reads that overlap the genomic origin for the peptide of interest are fetched from the RNA tumor BAM.

4. Reads are filtered to a reduced set containing the entirety of the peptide's coding sequence and counted. The resulting sum is provided as reads_with_peptide in the LENS report.

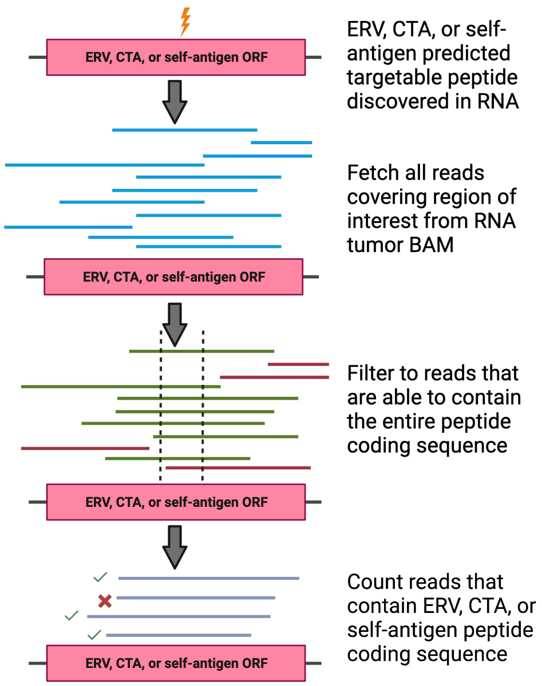

###### 4.1.5 Viruses

Oncogenic viruses are quantified in a manner similar the ERV, CTA, and self-antigen workflow as the entire viral protein can be considered for peptide generation. The primary difference is the read set considered when quantifying. Specifically,

1. The entirety of a detected viral protein is processed through NetMHCpan and other pMHC characterization tools.
2. Peptides of interest (e.g. high binding affinity) have their origin identified in the LENS viral reference.
3. Reads that *do not* map to the standard reference (e.g. hg38) but *do* map to the viral reference are considered.
4. Reads are filtered to a reduced set containing the entirety of the peptide's coding sequence and counted. The resulting sum is provided as reads_with_peptide in the LENS report.

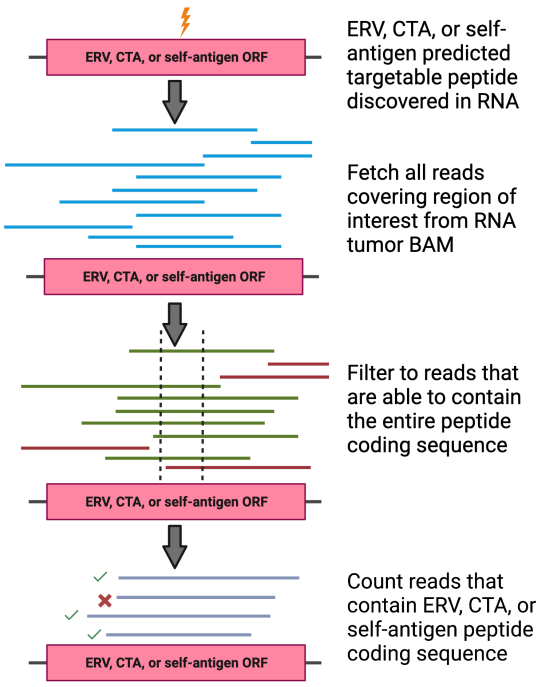

#### 4.2 Validating SNV and InDel Workflows with TESLA

The TESLA dataset allows for checking and benchmarking of the SNV and InDel workflows due to its set of experimentally validated immunogenic peptides.

### 5 Supplemental Tables

| Workflow | Inputs | MHC class support | Requires pre-processing | Workflow manager | Containerization | Harmonized Peptide Abundance Estimate | Modular (See caption) | HLA Typing | Peptide Manufacturability Considered | TCR repertoire | Somatic SNVs | Somatic in-frame InDels | Somatic frameshift InDels | Fusion Events | Splice Variants | Endogenous Retroviruses | Viruses | CTAs/ Self-Antigens |
|---|---|---|---|---|---|---|---|---|---|---|---|---|---|---|---|---|---|---|
| LENS | DNA tumor FASTQs<br>DNA normal FASTQs<br>RNA tumor FASTQs | Class I | ✗ | RAFT/Nextflow DSL2 | Process-level | ✓ | Supported through Nextflow DSL2 | ✓ | ✗ | ✓ | ✓ | ✓ | ✓ | ✓ | ✓ | ✓ | ✓ | ✓ |
| pVACtools | Annotated, phased VCFs (pVACseq)<br>Peptide FASTA (pVACbind)<br>Annotated Fusions (pVACfuse) | Class I & II | ✓ | N/A | ✗ | ✗ | ✗ | ✗ | ✓ | ✗ | ✓ | ✓ | ✓ | ✓ | ✗ | ✗ | ✗ | ✗ |
| OpenVax | DNA tumor FASTQs<br>DNA normal FASTQs<br>RNA tumor FASTQs | Class I | ✗ | SnakeMake<br>(optionally within Docker container) | Workflow-level | ✗ | Supported through SnakeMake | ✗ | ✓ | ✗ | ✓ | ✗ | ✗ | ✗ | ✗ | ✗ | ✗ | ✗ |
| nextNEOpi | DNA tumor FASTQs or BAM<br>DNA normal FASTQs or BAM<br>RNA tumor FASTQs or BAM (optional) | Class I & II | Optional<br>(short read alignment) | Nextflow DSL1 | Hybrid<br>(See caption) | ✗ | ✗ | ✓ | ✓ | ✓ | ✓ | ✓ | ✓ | ✓ | ✗ | ✗ | ✗ | ✗ |

**Supplemental Table 3: Comparison between LENS and other Neoantigen workflows:** LENS generally offers similar or improved functionality compared to other popular neoantigen workflows. Specifically, LENS supports more tumor antigen sources, does not require end-user pre-processing of input data, and is a both modular and extensible workflow. It is worth noting that all the included workflows are open-source and may subjectively be classified as modular and extensible as a result. We classify LENS as modular and extensible as these were desired features that drove design and development rather than consequences of code availability. The "Hybrid" classification for nextNEOpi's containerization metric is due to some components, like pVACtools and NeoFuse, having self-contained containers while other tools (e.g. variant callers) are all contained within a single container.

| Tumor Antigen Class | Reference Set | Filtering | General Workflow | Detection Method |
|---|---|---|---|---|
| SNVs | snpEff annotation:<br>"missense" | $>$ TPM 75th percentile &<br>$\geq$ 1 tumor RNA read supporting peptide CDS &<br>$<$ 500 nM binding affinity | BWA $\rightarrow$<br>Alignment Sanitization $\rightarrow$<br>Transcript Expression Filtering $\rightarrow$<br>Annotation $\rightarrow$<br>SNV Peptide Workflow | Detected in DNA;<br>Confirmed in RNA |
| InDels | snpEff annotation:<br>"disruptive insertion"<br>"disruptive deletion"<br>"conservative insertion"<br>"conservative deletion"<br>"frameshift" | $>$ TPM 75th percentile &<br>$\geq$ 1 tumor RNA read supporting peptide CDS &<br>$<$ 500 nM binding affinity | BWA $\rightarrow$<br>Alignment Sanitization $\rightarrow$<br>Transcript Expression Filtering $\rightarrow$<br>Annotation $\rightarrow$<br>InDel Peptide Workflow | Detected in DNA;<br>Confirmed in RNA |
| Splice Variants | NeoSplice detection with<br>GENCODE gtf and<br>tissue-matched normal sample | $>$ 20 k-mer expression in tumor &<br>$<$ 4 k-mer expression in normal &<br>$>$ 100 min. k-mer transcript coverage &<br>$\geq$ 1 tumor RNA read supporting peptide CDS &<br>$<$ 500 nM binding affinity | NeoSplice workflow $\rightarrow$<br>Splice Variant Peptide Workflow | Detected in RNA |
| Gene Fusions | STARFusion detection using CTAT Trinity library | $\geq$ 1 tumor RNA read supporting peptide CDS &<br>$<$ 500 nM binding affinity | STARFusion workflow $\rightarrow$<br>Fusion Peptide Workflow | Detected in RNA |
| Viruses | VirDetect reference | $>$ 1 reads mapped to viral sequence &<br>$\geq$ 25% RNA read coverage to viral sequence &<br>$\geq$ 1 tumor RNA read supporting peptide CDS &<br>$<$ 500 nM binding affinity | VirDetect Workflow$\rightarrow$<br>Viral Peptide Workflow | Detected in RNA |
| ERVs | gEVE database | $>$ 2-fold high expression in tumor than tissue-matched normal &<br>$\geq$ 50% RNA read depth to ORF sequence &<br>$\geq$ 1 tumor RNA read supporting peptide CDS &<br>$<$ 500 nM binding affinity | STAR $\rightarrow$<br>Salmon $\rightarrow$<br>edgeR differential expression $\rightarrow$<br>coverage filter $\rightarrow$<br>ERV Peptide Workflow | Detected in RNA |
| Cancer Self Antigens/<br>Self-antigen | CTDatabase CTA gene list | $>$ TPM 95th percentile &<br>$\geq$ 1 tumor RNA read supporting peptide CDS &<br>$<$ 500 nM binding affinity | STAR $\rightarrow$<br>Salmon $\rightarrow$<br>Transcript Expression Filtering $\rightarrow$<br>CTA/Self-antigen Peptide Workflow | Detected in RNA |

**Supplemental Table 4: Overview of Tumor Antigen Generation and Filtering:** LENS relies on external, publicly available references in order to generate tumor antigens. These references include somatic variant annotation tools (e.g. snpEff), fusion annotations (CTAT Trinity reference), as well as sequence-based references (VirDetect and gEVE). These references define the scope to which a peptide can be predicted – a peptide unable to be detected by the references cannot be detected by LENS. Tumor antigen target filtering is currently performed with liberal thresholds and filter refinement is a future goal.

| Function | Default Tool | Reason | Supported Alternatives | Reference |
|---|---|---|---|---|
| RNA Alignment | STAR | Reliable general-purpose aligner with good performance | bbmap | Baruzzo *et al.* (2017) |
| DNA Alignment | BWA | General-purpose aligner used in medical context | bbmap | Barbitoff *et al.* (2022) |
| InDel Realignment | ABRA2 | Coupled with Cadabra variant calling. IndelRealigner support dropped in GATK4 | none | Mose *et al.* (2019) |
| Somatic Variant Calling | MuTect2, Strelka2, Abra2 | Joint somatic callers | VarScan2 | Mose *et al.* (2019) |
| Germline Variant Calling | DeepVariant | Best performance and highest robustness among germline callers in Barbitoff *et al.* (2022) | GATK HaplotypeCaller | Barbitoff *et al.* (2022) |
| Gene Fusion Detection | STAR-Fusion | Good performance relative to other options | Arriba | Haas *et al.* (2019) |
| Variant Annotation | snpEff | Higher annotation concordance to HGVS nomenclature relative to ANNOVAR | none | Park and Park (2022) |
| Splice Variant Detection | NeoSplice | Highest sensitive among tested algorithms with simulated data | SplAdder | Chai *et al.* (2022b) |
| Duplicate Read Marker | Picard2 | Commonly used and supported duplicate read marker. Suports marking of inter-chromosomal duplicates. | samtools rmdup | http://broadinstitute.github.io/picard/ |
| Base Quality Recalibration | GATK4 | Only tool available for task to best of author's knowledge | none | N/A |
| Variant Phasing | WhatsHap | Well-supported and fast, but may be elevated switch error rate than competitors. Further benchmarking needed. | HaplotypeTools | Martin *et al.* (2016) |
| MHC Typing | arcasHLA | Good performance and support for both class I and class II alleles | HLAProfiler, Optitype, seq2HLA | Orenbuch *et al.* (2020) |

**Supplemental Table 5: Default Tool Decisions:** LENS is designed to be as flexible as possible with respect to tools and methods, but default tools are used. This table describes what tools are used, other supported alternatives, and the reason for picking each tool as the default.

| Tool/Workflow | LENS advantages | Days since last update* (Date of last update) | Notes |
|---|---|---|---|
| nextNEOpi | • Developed in Nextflow DSL2 instead of DSL1<br>• Supports more potential tumor antigen sources<br>• Modularity allows for tool swapping where appropriate<br>• More pMHC characterization tools supported<br>• Murine support | 220 days (August 3rd, 2022) | nextNEOpi is a workflow utilizing Nextflow DSL1 to allow an end-to-end experience for the user. Somatic peptide prediction is handled by pVACseq while fusion peptides are called by NeoFuse (a wrapper for Arriba and netMHCpan). |
| antigen.garnish | • End-to-end workflow for improved process tracking<br>• Modular implementation allows for newer version of NetMHC tools.<br>• Provides expression metric for predicted tumor antigens. | 248 days (July 6th, 2022) | antigen.garnish is utilized in LENS for foreignness and dissimilarity estimates. LENS fills a different role as it starts with raw data to generate reports while antigen.garnish assumes peptides have been already predicted by upstream tools. |
| CloudNeo | • End-to-end workflow for improved process tracking<br>• Developed in Nextflow DSL2 which allows LENS to run on AWS, GCP, or local clusters<br>• Covers more tumor antigen sources<br>• More and newer pMHC characterization tools supported | 1,422 days (April 19th, 2019) | CloudNeo is a neoantigen workflow developed in CWL Draft2 for running on the SevenBridges (currently Velsara) CGC platform. Users can manually run the workflow using Rabix. The workflow has been largely unmaintained. |
| DeepHLAPan | • End-to-end workflow for improved process tracking<br>• Developed in Nextflow DSL2 which allows LENS to run on AWS, GCP, or local clusters<br>• Includes expression information when summarizing pMHCS | 243 days (July 11th, 2022) | DeepHLAPan is a deep learning pMHC presentation and immunogenicity prediction tool. It is dissimilar from LENS in that it requires users to provide sample alleles and peptides rather than providing an end-to-end solution. DeepHLAPan is support within LENS as an pMHC characterization tool. |

Continued on next page. *Publicly available online repositories checked on March 11th, 2023.

| Tool/Workflow | LENS advantages | Days since last update* (Date of last update) | Notes |
|---|---|---|---|
| Epidisco | • Developed in Nextflow DSL2 which allows LENS to run on AWS, GCP, or local clusters<br>• Supports more potential tumor antigen sources<br>• Modularity allows for tool swapping where appropriate<br>• Supports running multiple patients simultaneously | 1,943 days (November 14th, 2017) | Epidisco is an early version of the PGV (Personalized Genomic Vaccine) workflow used in the PGV-001 clinical trial. It utilizes many of the same tools used in the later OpenVax workflow (including isovar and Vaxrank |
| Epi-seq | • Developed in Nextflow DSL2 which allows LENS to run on AWS, GCP, or local clusters<br>• Supports more potential tumor antigen sources<br>• Modularity allows for tool swapping where appropriate<br>• More pMHC characterization tools supported | 3,658 days (March 5th, 2013) | Epi-seq is an early neoantigen workflow which accepts a single set of RNA-Seq reads, calls tumor-specific SNVs using SNVQ, phases SNVs, and then predicts binding using 'NetMHC 3.0'. |
| INTEGRATE-Neo | • Developed in Nextflow DSL2 which allows LENS to run on AWS, GCP, or local clusters<br>• Supports more potential tumor antigen sources<br>• Modularity allows for tool swapping where appropriate<br>• More pMHC characterization tools supported | 1,845 days (February 20th, 2018) | INTEGRATE-Neo is a modifiied version of the INTEGRATE tool which not only detects gene fusions, but also extracts potential tumor antigens from fusions. The resulting fusion and peptide data are inputs into pVACFuse (part of the pVACtools suite). |
| MuPeXi | • End-to-end workflow for improved process tracking<br>• Developed in Nextflow DSL2 which allows LENS to run on AWS, GCP, or local clusters<br>• Supports more potential tumor antigen sources<br>• Modularity allows for tool swapping where appropriate<br>• More pMHC characterization tools supported | 1,495 days (February 5th, 2019) | MuPeXi is a workflow which, provided a VCF, set of HLA alleles, and expression TSV, predicts somatically-derived tumor antigens. It incorporates expression, but only at the gene-level (rather than transcript or allele-specific). |

Continued on next page. *Publicly available online repositories checked on March 11th, 2023.

| Tool/Workflow | LENS advantages | Days since last update* (Date of last update) | Notes |
|---|---|---|---|
| neoantimon | • End-to-end workflow for improved process tracking<br>• Developed in Nextflow DSL2 which allows LENS to run on AWS, GCP, or local clusters<br>• Supports more potential tumor antigen sources<br>• Modularity allows for tool swapping where appropriate<br>• More pMHC characterization tools supported<br>• Supports running multiple patients simultaneously | 221 days (August 2nd, 2022) | neoantimon is an R workflow designed for detecting tumor antigens from SNVs, InDels, and fusion structural variants. It requires users to provided an annotated VCF and patient-specific alleles. Optionally, users may provided RNA expression data, an RNA BAM, and CNV data to improve allele-specific expression and CCF estimates. |
| NeoANT-HILL | • Developed in Nextflow DSL2 which allows LENS to run on AWS, GCP, or local clusters<br>• Supports more potential tumor antigen sources<br>• Modularity allows for tool swapping where appropriate<br>• More pMHC characterization tools supported<br>• GRCh38 human genomes supported<br>• Murine support | 1,065 days (April 10th, 2020) | NeoANT-HILL is a workflow expecting either an annotated VCF and RNA-Seq data or simply RNA-Seq data. HLA calling is performed with Optitype. Only the GRCh37 reference is supported by the current release. |
| NeoFlow | • Developed in Nextflow DSL2 which allows LENS to run on AWS, GCP, or local clusters<br>• Supports more potential tumor antigen sources<br>• Modularity allows for tool swapping where appropriate<br>• More pMHC characterization tools supported | 302 days (May 13th, 2022) | Despite LENS advantages, NeoFlow is perhaps the most compelling and unique workflow among those currently available. NeoFlow allows processing of mass spectrometry data. This allows the leap from predicted tumor pMHCs to observable tumor pMHCs. NeoFlow is still limited to somatic variants for the time being, but the workflow serves as an inspiration to potential functional expansion within LENS. |
| neopepsee | • Developed in Nextflow DSL2 which allows LENS to run on AWS, GCP, or local clusters<br>• Supports more potential tumor antigen sources<br>• Modularity allows for tool swapping where appropriate<br>• More pMHC characterization tools supported | 1,650 days (September 3rd, 2018) | neopepsee is requires RNA-Seq FASTQs, a VCF of somatic variants, and, optionally, HLA alelles. This workflow reports neoantigens derived from somatic variants and includes relevant expression data in the form of TPMs. |

Continued on next page.*Publicly available online repositories checked on March 11th, 2023.

| Tool/Workflow | LENS advantages | Days since last update* (Date of last update) | Notes |
|---|---|---|---|
| NeoPredPipe | • Developed in Nextflow DSL2 which allows LENS to run on AWS, GCP, or local clusters<br>• Supports more potential tumor antigen sources<br>• Modularity allows for tool swapping where appropriate<br>• More pMHC characterization tools supported | 716 days (March 25th, 2021) | NeoPredPipe requires an input VCF containing somatic variants, HLA alleles, and optionally, RNA-Sequencing data. A unique aspect of NeoPredPipe is the ability to consider variants called from multiple bioposies from a single patient. This information provides crucial information regarding the heterogeneity of the tumor and allows NeoPredPipe's outputs to applied to the neoantigen fitness model from Łuksza *et al.* (2017). |
| NeoepitopdePred | • Developed in Nextflow DSL2 which allows LENS to run on AWS, GCP, or local clusters<br>• Supports more potential tumor antigen sources<br>• Modularity allows for tool swapping where appropriate<br>• More pMHC characterization tools supported | 1,928 days* (November 29th, 2017) | NeoepitopePred is a somatic missense variant a fusion tumor antigen workflow which requires DNANexus cloud support for running.<br>* The original repository containing NeoepitopdePred no longer exists and the workflow is not listed under supported software for the lab that originally developed it. This push references a repository under GitHub user XingTang2014. |
| Neoepiscope | • Developed in Nextflow DSL2 which allows LENS to run on AWS, GCP, or local clusters<br>• Supports more potential tumor antigen sources<br>• Modularity allows for tool swapping where appropriate | 222 days (August 1st, 2022) | Neoepiscope is workflow for predicting somatic missense and indel-derived peptides neoantigens. This workflow puts a very heavy emphasis on phasing of potentially targetable variants with their surrounding genomic context in order to maximize sequence accuracy during peptide prediction. LENS also attempt to achieve this level of accuracy by phasing germline and somatic variants around targetable variants of interest. |
| NeoFuse | • Developed in Nextflow DSL2 which allows LENS to run on AWS, GCP, or local clusters<br>• Supports more potential tumor antigen sources<br>• Modularity allows for tool swapping where appropriate | 214 days (August 9th, 2022) | NeoFuse combines Arriba for fusion detection, Optitype from HLA typing, andNetMHCpan for pMHC charactierization to discover gene fusion-derived tumor antigens. It serves as the fusion caller within the nextNEOpi workflow. |

Continued on next page.*Publicly available online repositories checked on March 11th, 2023.

| Tool/Workflow | LENS advantages | Days since last update* (Date of last update) | Notes |
|---|---|---|---|
| OpenVax | • End-to-end workflow for improved process tracking<br>• Supports more potential tumor antigen sources<br>• Modularity allows for tool swapping where appropriate | 933 days (August 20th, 2020) | The neoantigen-vaccine-pipeline, released by the OpenVax team, is the spiritual successor to the 'Epidisco' workflow described above. Overall, the workflow is very similar to LENS and introduces a unique form of variant phasing using RNA sequencing reads. It utilizes snakemake for workflow management and 'Docker' to allow for easier dependency resolution for end users. |
| ProGeo-Neo | • End-to-end workflow for improved process tracking<br>• Supports more potential tumor antigen sources<br>• Modularity allows for tool swapping where appropriate | 339 days (April 6th, 2022) | The ProGeo-Neo is a workflow requiring user-provided RNA-Seq reads, HLA types, a sample-specific protein database, and mass spectroscopy data. Users must provide all workflow dependencies which may hamper reproducibility, but the utilization of mass spectroscopy data makes this workflow unique. |
| pTuneos | • Developed in Nextflow DSL2 which allows LENS to run on AWS, GCP, or local clusters<br>• Supports more potential tumor antigen sources<br>• Modularity allows for tool swapping where appropriate | 482 days (November 14th, 2021) | pTuneos is an end-to-end SNV/InDel neoantigen workflow which utilizes a machine learning-based approach to predict pMHC immunogenecity. |
| pVACtools | • Developed in Nextflow DSL2 which allows LENS to run on AWS, GCP, or local clusters<br>• Supports more potential tumor antigen sources<br>• Modularity allows for tool swapping where appropriate | 52 days (Janurary 18th, 2023) | The pVACtools suite is perhaps one of the best known neoantigen workflows under active development. It includes utilities to predict SNV-, InDel-, and fusion-derived pMHCs and includes usefuls utilities for guiding vaccine development. |
| nfcore-epitopeprediction | • End-to-end workflow for improved process tracking Modularity allows for tool swapping where appropriate | 8 days (March 3rd, 2023) | nfcore-epitopeprediction is a component of the nfcore suite of Nextflow workflows. It requires that users provide VCFs and alleles in order to predict peptides and process them through a variety of pMHC characterization tools. |

Continued on next page. *Publicly available online repositories checked on Match 11th, 2023.

| Tool/Workflow | LENS advantages | Days since last update* (Date of last update) | Notes |
|---|---|---|---|
| ScanNeo | • Developed in Nextflow DSL2 which allows LENS to run on AWS, GCP, or local clusters<br>• Supports more potential tumor antigen sources<br>• Modularity allows for tool swapping where appropriate<br>• Does not require manually modified hardcoded reference file paths | 194 days (August 29th, 2022) | The ScanNeo is a workflow designed to detect Class I SNV- and InDel-derived tumor antigens with both human and murine support. |
| TIminer | • Developed in Nextflow DSL2 which allows LENS to run on AWS, GCP, or local clusters<br>• Supports more potential tumor antigen sources<br>• Modularity allows for tool swapping where appropriate<br>• Does not require manually modified hardcoded reference file paths | 2,280 days (December 12th, 2016) | TIminer is a workflow for discovering SNV/InDel class I tumor antigens. It is able to perform gene set enrichment analysis (GSEA) in order to inform peptide filtering with an "immunophenogram" and "immunophenoscore". It also includes a user interface for setting up workflows which improves the end-user experience. |
| TSNAD | • Developed in Nextflow DSL2 which allows LENS to run on AWS, GCP, or local clusters<br>• Supports more potential tumor antigen sources<br>• Modularity allows for tool swapping where appropriate | 578 days (August 20th, 2021) | TSNAD is a dockerized end-to-end Class I SNV/InDel and fusion neoantigen workflow from the authors of DeepHLAPan. |
| VaxRank | • Developed in Nextflow DSL2 which allows LENS to run on AWS, GCP, or local clusters<br>• Supports more potential tumor antigen sources<br>• Modularity allows for tool swapping where appropriate | 934 days (August 19th, 2020) | VaxRank is a tool for calculating and prioritizing immunogencity from a user-provided VCF and BAM file. It supports a variety of pMHC characterization tools. It is a crucial component of the Neoantigen-Vaccine-Pipeline by the OpenVax team. |
| TruNeo | • Developed in Nextflow DSL2 which allows LENS to run on AWS, GCP, or local clusters<br>• Supports more potential tumor antigen sources<br>• Modularity allows for tool swapping where appropriate | 836 days (November 25th, 2020) | VaxRank is a tool for calculating and prioritizing immunogencity from a user-provided VCF and BAM file. It supports a variety of pMHC characterization tools. It is a crucial component of the Neoantigen-Vaccine-Pipeline by the OpenVax team. |

*Publicly available online repositories checked on March 11th, 2023.

**Supplemental Table 6: LENS Compared to Competition:** There are a variety of neoantigen workflows available. Here we have compared LENS to over two dozen other workflows.

### References

(2022). Endogenous retroviruses in acute myeloid leukemia (erval). https://clinicaltrials.gov/ct2/show/NCT04406207. Accessed: 2022-03-22.

An, X. *et al.* (2014). Global transcriptome analyses of human and murine terminal erythroid differentiation. *Blood, The Journal of the American Society of Hematology*, **123**(22), 3466–3477.

Anguille, S. *et al.* (2012). Leukemia-associated antigens and their relevance to the immunotherapy of acute myeloid leukemia. *Leukemia*, **26**(10), 2186–2196.

Barbitoff, Y. A. *et al.* (2022). Systematic benchmark of state-of-the-art variant calling pipelines identifies major factors affecting accuracy of coding sequence variant discovery. *BMC genomics*, **23**(1), 155.

Baruzzo, G. *et al.* (2017). Simulation-based comprehensive benchmarking of rna-seq aligners. *Nature methods*, **14**(2), 135–139.

Cellier, M. F. (2013). Cell-type specific determinants of nramp1 expression in professional phagocytes. *Biology*, **2**(1), 233–283.

Chai, S. *et al.* (2022a). NeoSplice: a bioinformatics method for prediction of splice variant neoantigens. *Bioinformatics Advances*, **2**(1).

Chai, S. *et al.* (2022b). Neosplice: a bioinformatics method for prediction of splice variant neoantigens. *Bioinformatics Advances*, **2**(1), vbac032.

Chen, G. *et al.* (2005). Serological identification of immunogenic antigens in acute monocytic leukemia. *Leukemia research*, **29**(5), 503–509.

Chen, X. *et al.* (2021). Fusion gene map of acute leukemia revealed by transcriptome sequencing of a consecutive cohort of 1000 cases in a single center. *Blood Cancer Journal*, **11**(6), 1–10.

Gilman, P. *et al.* (2019). Pysam (python wrapper for system advisor model" sam"). Technical report, National Renewable Energy Lab.(NREL), Golden, CO (United States).

Greif, P. A. *et al.* (2011). Identification of recurring tumor-specific somatic mutations in acute myeloid leukemia by transcriptome sequencing. *Leukemia*, **25**(5), 821–827.

Haas, B. J. *et al.* (2019). Accuracy assessment of fusion transcript detection via read-mapping and de novo fusion transcript assembly-based methods. *Genome biology*, **20**(1), 1–16.

Kandoth, C. *et al.* (2013). Mutational landscape and significance across 12 major cancer types. *Nature*, **502**(7471), 333–339.

Lee, J. H. *et al.* (2016). A novel kit indel mutation in acute myeloid leukemia with t(8;21)(q22;q22); runx1-runx1t1. *Annals of Laboratory Medicine*, **36**(4), 371–374.

Li, S. *et al.* (2019). Mpo as a novel susceptibility gene in myeloid malignancies. *Blood*, **134**, 5402.

López-Nieva, P. *et al.* (2019). Detection of novel fusion-transcripts by rna-seq in t-cell lymphoblastic lymphoma. *Scientific reports*, **9**(1), 1–11.

Łuksza, M. *et al.* (2017). A neoantigen fitness model predicts tumour response to checkpoint blockade immunotherapy. *Nature*, **551**(7681), 517–520.

Martin, M. *et al.* (2016). Whatshap: fast and accurate read-based phasing. *BioRxiv*, page 085050.

Masson, D. *et al.* (1986). Identification of granzyme a isolated from cytotoxic t-lymphocyte-granules as one of the proteases encoded by ctl-specific genes. *FEBS letters*, **208**(1), 84–88.

Mose, L. E. *et al.* (2019). Improved indel detection in dna and rna via realignment with abra2. *Bioinformatics*, **35**(17), 2966–2973.

Nakagawa, S. and Takahashi, M. U. (2016). geve: a genome-based endogenous viral element database provides comprehensive viral protein-coding sequences in mammalian genomes. *Database*, **2016**.

Orenbuch, R. *et al.* (2020). arcashla: high-resolution hla typing from rnaseq. *Bioinformatics*, **36**(1), 33–40.

Park, K.-J. and Park, J.-H. (2022). Variations in nomenclature of clinical variants between annotation tools. *Laboratory Medicine*, **53**(3), 242–245.

Quintarelli, C. *et al.* (2011). High-avidity cytotoxic t lymphocytes specific for a new prame-derived peptide can target leukemic and leukemic-precursor cells. *Blood, The Journal of the American Society of Hematology*, **117**(12), 3353–3362.

She, J. *et al.* (2022). The landscape of hervrnas transcribed from human endogenous retroviruses across human body sites. *Genome Biology*, **23**(1), 1–21.

Steinbach, D. *et al.* (2006). Identification of a set of seven genes for the monitoring of minimal residual disease in pediatric acute myeloid leukemia. *Clinical cancer research*, **12**(8), 2434–2441.

van der Lee, D. I. *et al.* (2019). Mutated nucleophosmin 1 as immunotherapy target in acute myeloid leukemia. *The Journal of clinical investigation*, **129**(2), 774–785.

van der Lee, D. I. *et al.* (2021). An hla-a* 11: 01-binding neoantigen from mutated npm1 as target for tcr gene therapy in aml. *Cancers*, **13**(21), 5390.

Zhou, J. and Chng, W.-J. (2017). Aberrant rna splicing and mutations in spliceosome complex in acute myeloid leukemia. *Stem Cell Investigation*, **4**, 6.

Zhou, J. X. *et al.* (2017). Identification of kansarl as the first cancer predisposition fusion gene specific to the population of european ancestry origin. *Oncotarget*, **8**(31), 50594.
